# CD38 promotes hematopoietic stem cell dormancy via c-Fos

**DOI:** 10.1101/2023.02.08.527614

**Authors:** Liliia Ibneeva, Sumeet Pal Singh, Anupam Sinha, Sema Elif Eski, Rebekka Wehner, Luise Rupp, Juan Alberto Pérez-Valencia, Alexander Gerbaulet, Susanne Reinhardt, Manja Wobus, Malte von Bonin, Jaime Sancho, Frances Lund, Andreas Dahl, Marc Schmitz, Martin Bornhäuser, Triantafyllos Chavakis, Ben Wielockx, Tatyana Grinenko

## Abstract

A subpopulation of deeply quiescent, so-called dormant hematopoietic stem cells (dHSCs) resides at the top of the hematopoietic hierarchy and serves as a reserve pool for HSCs possessing the greatest long-term blood repopulation capacity. The state of dormancy protects the HSC pool from exhaustion throughout life, however excessive dormancy may block an efficient response to hematological stresses. The mechanisms of HSC dormancy remain elusive, mainly due to the absence of surface markers that allow dHSC prompt isolation. Here, we identify CD38 as a novel surface marker for murine dHSCs that is broadly applicable. Moreover, we demonstrate that cyclic adenosine diphosphate ribose (cADPR), the product of CD38 cyclase activity, regulates the expression of the transcription factor c-Fos by increasing cytoplasmic Ca^2+^ concentration. Strikingly, we uncover that c-Fos drives HSCs dormancy through the induction of the cell cycle inhibitor p57^Kip2^. Moreover, we found that CD38 ecto-enzymatic activity at the neighboring CD38-positive cells can promote human HSC quiescence. Together, CD38/cADPR/Ca^2+^/cFos/p57^Kip2^ axis maintains HSC dormancy. Pharmacological manipulations of this pathway can provide new strategies to expand dHSCs for transplantation or to activate them during hematological stresses.

## Introduction

Hematopoietic stem cells (HSCs) are responsible for the production of all blood cells during life. Adult HSCs are maintained in a quiescent state, which is thought to protect them not only from replicative and metabolic stresses but also from the accumulation of somatic mutations; thus, loss of quiescence could lead to their exhaustion or malignant transformation (1–4). Conversely, excessive quiescence can lead to the generation of too few blood cells, which can result in reduced immune responses and greater infection susceptibility. Therefore, tight regulation of the balance between HSC quiescence and activation is critical for effective hematopoiesis under normal and stress conditions.

Numerous studies have demonstrated that 20–30% of murine HSCs are deeply quiescent, that they do not produce cells under homeostatic conditions, and that these ‘dormant’ HSCs (dHSCs) (4) harbor the greatest long-term repopulation capacity in transplantation assays (4–6). Thus, dHSCs serve as a reserve pool of stem cells that are activated only in response to stress signals such as interferons, lipopolysaccharide, and myeloablation, thereby demonstrating their importance in organismal stress resistance and recovery after chemotherapy (4, 7). However, despite the importance of mechanisms that switch HSCs from dormant to active state, detailed characterization of such dHSCs has been challenging due to the absence of known surface markers for their ready identification and isolation. Consequently, processes involved in the preservation of dHSC quiescence are poorly understood.

Recently, Cabezas-Wallscheid et al., have established a Gprc5c (retinol receptor) reporter mouse strain and have shown that retinoic acid signaling and hyaluronic acid could regulate HSC dormancy (6, 8). Fukushima et al., used another p27 (Cdk inhibitor) reporter mouse strain to reveal that HSC entry in to the cell cycle is controlled by Cdk4/6 and that high cytosolic Ca^2+^ concentration correlates with HSC quiescence (9). However, why dHSCs harbor high cytosolic Ca^2+^ concentration and how Ca^2+^ regulates HSC dormancy remain unclear. Here, we identify that CD38 is not only the surface marker for the isolation of murine dHSCs but also describe a previously unknown signaling axis driven by the ecto-enzymatic activity of CD38 controlling HSC dormancy. Mechanistically, we show that cyclic adenosine diphosphate ribose (cADPR), the product of nicotinamide adenine dinucleotide (NAD) conversion by CD38, regulates the expression of the transcription factor c-Fos, thereby driving quiescence in a p57^Kip2^-dependent manner.

## Results

### Pseudotime analysis of HSCs reveals the transition between proliferation and dormancy

To capture the transition between quiescence and proliferation, HSCs from young mice were subjected to single cell RNA sequencing, and after quality control, the transcriptome profiles of 1613 individual HSCs were used for downstream analysis (Fig. 1A). To identify actively cycling cells, we calculated cell cycle and dormancy scores of individual HSCs using Seurat (10), which were based on the expression levels of cell cycle and dormancy genes (11) (Suppl. Table 1). We observed that cells in the S (Fig. 1B) and the G2/M phases (Fig. 1C) were clustered together and that, as expected, most of the HSCs were quiescent (Fig. 1D). Further, comparison of Fig. 1A with Fig. 1B–D showed that pseudotime ordering was congruent with the transition from proliferation to dormancy. Next, we applied pseudotime ordering (Fig. 1A) to identify gene expression patterns that correlate with cell cycle dynamics (Fig. 1E) and identified three major gene expression clusters, namely, 1 - Early, 2 - Intermediate, and 3 - Late genes, according to peak expression in relation to pseudotime (Fig. 1E, Suppl. Table 2). Functional annotation of each cluster revealed that Early and Intermediate genes were typically related to cell cycle activation pathways (Fig. 1F, Supp. Table 3). In contrast, Late genes included well-known markers of HSCs with high transplantation potential, i.e., *Vwf* (12), *Procr* (13), *Fgd5* (14, 15), and the cell cycle inhibitor *Cdkn1c* (16, 17) (Fig. 1G, Fig. S1 A, B). Notably, this cluster was characterized by genes involved in pathways related to the activation of tumor necrosis factor alpha (TNFα) signaling, interferon gamma and alpha response, Stat3 and Stat5, as well as transforming growth factor beta 1 (TGF-β1) signaling, which is a well-known regulator of HCS quiescence (18) (Fig. 1F, Suppl. Table 3). Next, we attempted to isolate cell surface markers within the group of Late genes (Suppl. Table 2) associated with HSC dormancy and identified *Cd38* as a putative marker for dHSCs because its expression was higher in cells with high dormancy scores and corresponded with expression of well-known long-term HSC (LT-HSC) markers (Fig. 1 G, H).

**Figure 1.**
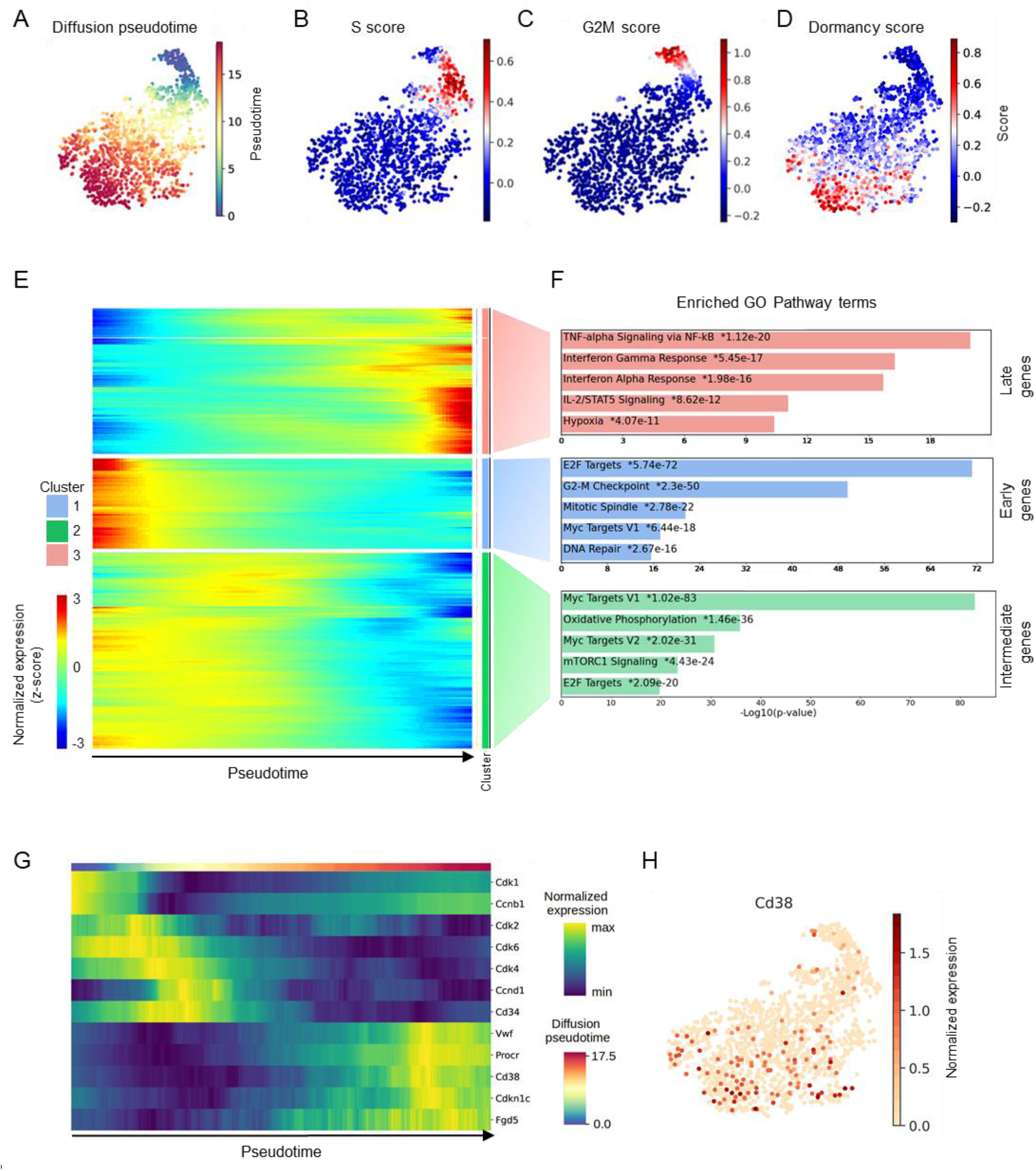
Single cell transcriptome analysis of HSCs. (A) Uniform manifold approximation projection (UMAP) representation depicting the transcriptional profiles of individual HSCs (LSK CD48^-^ CD150^+^). (B) S-phase score along pseudotime. (C) G2/M phase score along pseudotime. (D) Dormancy score along pseudotime. For panels A-D, each dot represents a single cell. (E) Clustered heatmap showing expression of genes along pseudotime. Each column represents a single cell and each row represents a single gene. (F) The first five most-significantly enriched pathways in each cluster are shown. (G) Expression of selected genes along pseudotime. (H) UMAP representation showing the expression of *Cd38*. Each dot represents a single cell.

### CD38^+^ LT-HSCs harbor the highest repopulation capacity

We analyzed surface expression of CD38 on hematopoietic stem and progenitor cells (HSPCs), and showed that CD38 was expressed by fractions of LT-HSCs (Lin^-^ c-Kit^+^ Sca-1^+^ (LSK) CD48^-^ CD150^+^ CD34^-^ CD201^+^; 36.6 ± 2.5%), HSCs (LSK CD48^-^ CD150^+^; 12.4 ± 0.7%) and multipotent progenitors 2 (MPP2, LSK CD48^+^ CD150^+^; 15.3 ± 1.8%) but not short-term HSCs (ST-HSCs, LSK CD48^-^ CD150^-^) or multipotent progenitors 3/4 (MPP3/4, LSK CD48^+^ CD150^-^) (Fig. 2A-B, Fig. S2 A, B). Next, we subdivided HSCs based on CD38 surface expression as CD38^+^ and CD38^-^ stem cells and compared the expression of well-known surface markers defining the most potent LT-HSCs (4, 19–23), and revealed that, compared to CD38^-^ HSCs, surface expression of CD34, CD229, and c-Kit were lower, while that of CD201, Sca-1, CD150 were higher in CD38^+^ HSCs (Fig. S2C). In line with these data, the frequency of LT-HSCs was higher among CD38^+^ HSCs compared to other fractions of the HSCs (Fig. 2C). Together, these data indicate that CD38^+^ HSCs display a phenotype identical to that of the most potent and quiescent LT-HSCs (4, 19–23).

**Figure 2.**
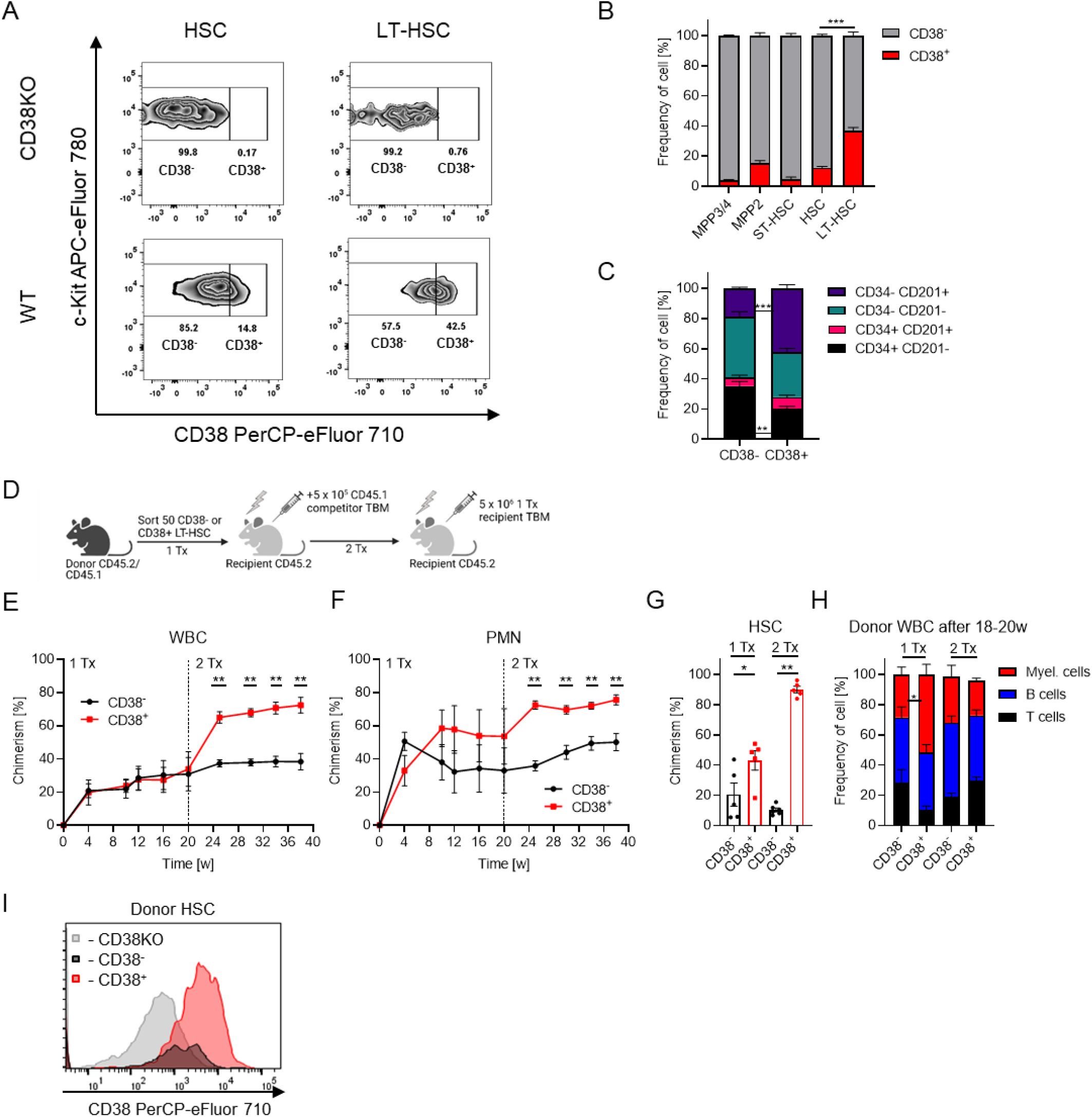
CD38^+^ defines LT-HSCs with the highest long-term repopulation capacity. (A) Flow cytometry analysis of CD38 expression on HSCs (LSK CD48^-^ CD150^+^) and LT-HSCs (LSK CD48^-^ CD150^+^ CD34^-^ CD201^+^) in CD38KO (negative control for staining) vs wt mice. (B) Frequency of CD38^+^ cells in various HSPC populations, n=7. Multiple-group comparisons were performed using Brown-Forsythe and Welch ANOVA followed by Dunnett’s T3 multiple comparison tests. ***p<0.001. (C) Frequencies of different HSC subpopulations in CD38- and CD38^+^ HSCs, n=7. (D) Set-up for CD38^-^ and CD38^+^ LT-HSCs (LSK CD48^-^ CD150^+^ CD201^+^ CD34^-^) transplantation, 2 independent experiments, 1 representative experiment is shown, n=5. (E) Chimerism in donor-derived WBC cells after transplantation. (F) Chimerism in donor-derived polymorphonuclear neutrophils (PMN) after transplantation, PMN: Gr1 ^+^ CD11b^+^. (G) Chimerism in the HSC population after transplantation. (H) Frequency of T, B, and myeloid cells in donor derived peripheral blood (PB) cells at 18-20 weeks after transplantation. (I) Surface expression of CD38 in donor-derived HSCs at 20 weeks after primary transplantation of CD38^+^ or CD38^-^ LT-HSCs (5 mice pooled), CD38 knock-out HSCs were used as negative control for CD38 staining. C, E-H - p-value was calculated using Mann-Whitney *U*-test, *p<0.05, **p<0.01.

Only a fraction of LT-HSCs ((LSK) CD48^-^ CD150^+^ CD34^-^ CD201^+^) expresses CD38. To assess whether CD38 expression correlates with the superior repopulation capacity within LT-HSCs, we transplanted CD38^+^ and CD38^-^ LT-HSCs into lethally irradiated mice under competitive settings (Fig. 2D). While CD38^-^ LT-HSCs produced more short-lived neutrophils four weeks after transplantation, CD38^+^ LT-HSCs repopulated the HSC compartment and peripheral blood (PB) more efficiently at 20 weeks after primary transplantation and after secondary transplantation as well (Fig. 2 E–G). Further, no lineage bias was observed in the reconstitution pattern of CD38^+^ and CD38^-^ LT-HSCs (Fig. 2H). These results demonstrate the superior repopulation and self-renewal capacity of CD38^+^ cells compared to CD38^-^ LT-HSCs.

To understand the hierarchy between CD38^+^ and CD38^-^ LT-HSCs, we compared the expression of CD38 on the progeny of donor HSCs and found that, while CD38^+^ LT-HSCs gave rise to both CD38^-^ and CD38^+^ HSCs, CD38^-^ cells could not generate CD38^+^ HSCs (Fig. 2I). Taken together, these results indicate that CD38^+^ LT-HSCs reside at the top of the hematopoietic cell hierarchy. We propose that the CD38 surface marker can be used atop to the well-established immuno-phenotype of LSK CD48^-^ CD150^+^ CD34^-^ CD201^+^ to define the most potent LT-HSCs.

### High levels of surface CD38 define dormant HSCs

We performed cell cycle analyses, bromodeoxyuridine (BrdU) incorporation, and long-term label-retaining assays to investigate whether CD38 expression correlates with stem cells’ dormancy (Fig. 3A-F). CD38^+^ HSCs mostly resided in G0 phase compared with CD38^-^ HSCs (Fig. 3A). While LT-HSC markers already enrich for more quiescent cells, CD38^+^ LT-HSCs contained an even higher frequency of cells in G0 phase than CD38^-^ LT-HSCs (Fig. 3B). Accordingly, CD38^+^ LT-HSCs incorporated BrdU significantly slower and retained higher levels of H2B-GFP after 130 days of chase compared with CD38^-^ cells (Fig. 3D-F), revealing that CD38^+^ LT-HSCs are more quiescent than their CD38^-^ counterparts (24). Moreover, CD38^+^ LT-HSCs had lower mitochondrial membrane potential (MMP) than CD38^-^ stem cells, despite no difference in mitochondrial mass (Fig. S2 D, E), which is in agreement with previous findings that HSC quiescence is associated with a lower metabolic status (6, 25).

**Figure 3.**
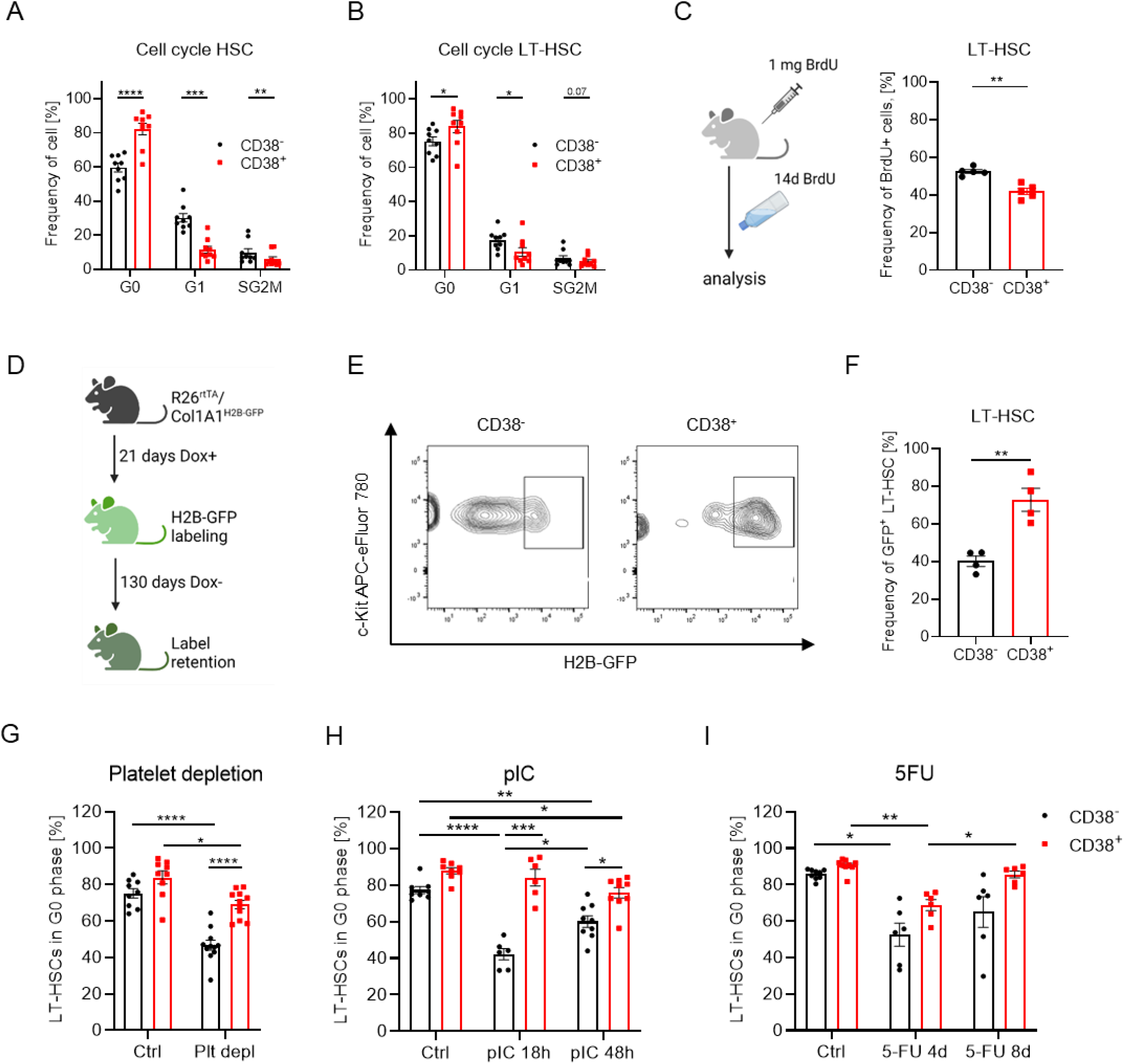
CD38^+^ dHSCs reside at the top of LT-HSC hierarchy. (A) Frequencies of CD38^-^ and CD38^+^ HSCs or LT-HSCs (B) in G0, G1 and SG2M phases of the cell cycle, n=9. (C) BrdU incorporation assay. Frequency of BrdU^+^ cells in CD38^-^ and CD38^+^ LT-HSC populations 14d after BrdU, n=5. (D) Set-up of a long-term label retention assay. (E) Gating strategy for the identification of GFP-retaining LT-HSCs. (F) Frequency of GFP^+^ cells in CD38^-^ and CD38^+^ LT-HSCs; n=4. (G) Cell cycle analysis of LT-HSCs at 18 h after platelet depletion (n=10-11; 3 independent experiments). (H) Cell cycle analysis of LT-HSCs at 18 and 48 h after pIC injection (n=6-9; 3 independent experiments). (I) Cell cycle analysis of LT-HSCs at days 4 and 8 after 5-FU injection (n=6-11; 3 independent experiments). For panels A-C, F - the paired *t*-test was used. Multiple-group comparisons were performed using Brown-Forsythe and Welch ANOVA followed by Dunnett’s T3 multiple comparison tests. *p<0.05, **p< 0.01, ***p<0.001, ****p<0.0001.

Next, to investigate cell cycle entry of CD38^+^ LT-HSCs in response to hematopoietic stressors, mice were administered anti-platelet serum to mimic acute autoimmune thrombocytopenia (26), polyI:polyC (pIC) mimicking viral infection (6), or 5-fluorouracil (5-FU), a myeloablative agent. We found that while CD38^-^ LT-HSCs partially proliferated in response to platelet depletion, CD38^+^ LT-HSCs retained their quiescence (Fig. 3G). Likewise, while CD38^-^ LT-HSC rapidly entered the cell cycle in response to pIC, fewer CD38^+^ LT-HSCs entered the cell cycle and did so with a significant delay (Fig. 3H, S2F). In contrast, both CD38^-^ and CD38^+^ LT-HSCs actively proliferated 4 days after 5-FU injection (Fig. 3I). Although CD38^+^ LT-HSCs tended to restore their quiescence 8 days after 5-FU injection, CD38^-^ cells remained in the cell cycle. Thus, as CD38^+^ LT-HSCs required more time to enter the cell cycle in response to hematological stresses and returned faster to quiescence compared to CD38^-^ cells, we posit that high levels of CD38 expression define dormant LT-HSCs not only in steady state but also under hematopoietic stress. Taken together, we classified CD38^+^ LT-HSCs as dHSCs.

### CD38 enzymatic activity regulates dHSC quiescence

To understand whether CD38 is directly involved in the maintenance of dHSCs, we compared long-term repopulation and self-renewal capacities of wild-type (wt) and CD38 knock-out (CD38KO) LT-HSCs. We did not find any difference in composition of PB or bone morrow at steady state (Fig. S3 A–F). Although only about 40% of LT-HSC in wt mice express CD38, we found that long-term repopulation and self-renewal capacity of CD38KO LT-HSC were lower than those of wt cells (Fig. S3 G–K). In line with this finding, CD38KO total bone marrow (TBM) cells had diminished long-term repopulation capacity compared with wt TBM (Fig. S3L-P). Together, these results suggest that CD38 regulates the functionality of LT-HSCs.

CD38 is a multifaceted NAD catabolic ecto-enzyme that metabolizes NAD and its precursors (nicotinamide mononucleotide-NMN and nicotinamide riboside-NR) into adenosine diphosphate ribose (ADPR) and cyclic-ADPR (cADPR) (27) (Fig. 4A). 78c is a specific CD38 inhibitor that hinders both hydrolase and ADP-ribosyl cyclase activities of CD38 (28). To investigate whether the enzymatic activity of CD38 regulates the quiescence of LT-HSCs, we performed single-cell tracing experiments wherein LT-HSC division in the presence of 78c was tracked (Fig. S4 A–C, Fig. 4B-C). In agreement with our previous data (Fig. 3), CD38^+^ cells were more quiescent than CD38^-^ LT-HSCs, whereas inhibition of CD38 by 78c accelerated the first division of CD38^+^ but not CD38^-^ LT-HSCs or LT-HSCs from CD38KO mice (Fig. 4B-C, Fig. S4 A–C), supporting the idea that CD38 enzymatic activity indeed contributes to maintenance of dHSC quiescence.

**Figure 4.**
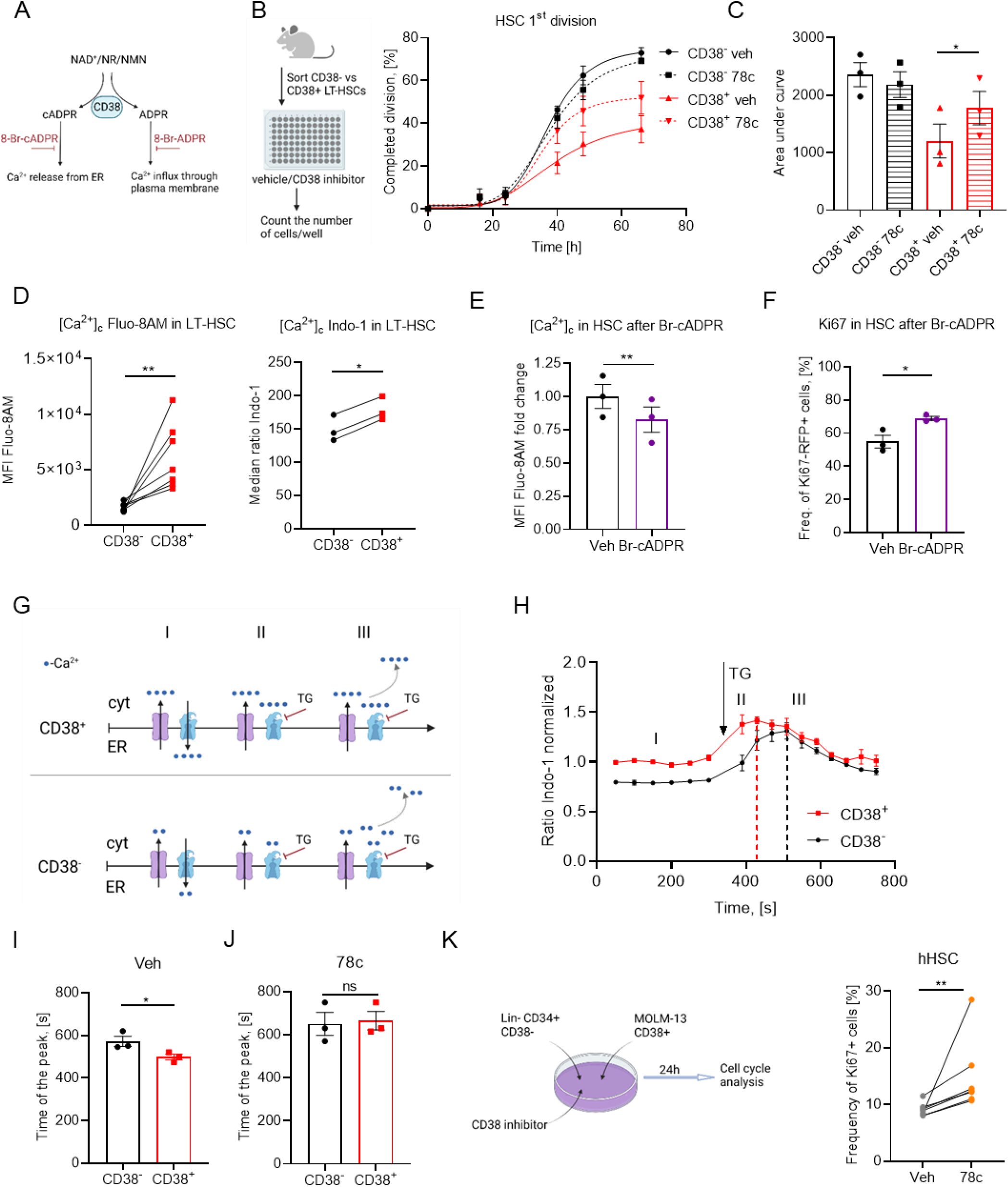
CD38 enzymatic activity regulates dHSC dormancy. (A) Enzymatic activities of CD38. (B) Single CD38^-^ and CD38^+^ LT-HSCs were sorted and cultured in liquid media with or without 78c, a CD38 inhibitor. Frequency of LT-HSCs that had completed the first division during incubation time is presented (3 independent experiments). (C) Quantification of AUC for data in panel B. (D) Free cytoplasmic Ca^2+^ ([Ca^2+^]_c_) relative concentration in CD38^+^ and CD38^-^ LT-HSCs analyzed using Fluo-8AM dye, n=7, and ratiometric Indo-1 dye, n=3. (E) Relative [Ca^2+^]_c_ concentration in HSCs treated with Br-cADPR for 24 h, n=3. 2 independent experiments. (F) Frequency of cycling in Ki67-RFP^+^ HSCs at 24 h after treatment with Br-cADPR, n=3, 2 independent experiments. (G) Suggested model of [Ca^2+^]_c_ modulation. Time frame I: Under steady state conditions, Ca^2+^ is released into the cytoplasm and pumped back into the ER. CD38^+^ HSCs release more Ca^2+^ from ER than CD38^-^ cells due to cADPR, the product of CD38 enzymatic activity. Time frame II: Blocking Ca^2+^-ATPase pumping Ca^2+^ into the ER with TG led to a faster rise in [Ca^2+^]_c_ concentration CD38^+^ HSCs than in CD38^-^ cells. Time frame III: excessive [Ca^2+^]_c_ is removed from cytoplasm. (H) [Ca^2+^]_c_ dynamics in CD38^-^ and CD38^+^ HSCs. (I) Time between addition of TG and maximum [Ca^2+^]_c_ in CD38^-^ and CD38^+^ HSCs, n=3. (J) Time between addition of TG and maximum [Ca^2+^]_c_ in CD38^-^ and CD38^+^ HSCs in the presence of a CD38 inhibitor, n=3. (K) Co-culture of hHSCs from healthy adult donors and MOLM-13 in the presence of CD38 inhibitor 78c during 24h. Cell cycle of hHSCs was analyzed (n=7). For panels C, D, F-J P-values were calculated using the paired *t*-test; for panel E, the unpaired *t*-test was used; for panel K, the Mann-Whitney *U*-test was used *p<0.05, **p<0.01.

Both products of CD38 enzymatic activity, i.e., ADPR and cADPR, increase cytosolic Ca^2+^ concentration ([Ca^2+^]_c_) in several cell types (29–31) (Fig.4A) and high cytoplasmic Ca^2+^ has been shown to support quiescence of HSCs (9). Consistently, we show that [Ca^2+^]_c_ is indeed higher in CD38^+^ LT-HSCs than in CD38^-^ cells when measured using Ca^2+^-indicator dyes, Fluo-8AM and Indo-1 (Fig. 4D). Additionally, while treatment of cells with 8-Bromo-ADPR (a cell-permeable antagonist that blocks ADPR-dependent Ca^2+^ release) did not influence either [Ca^2+^]_c_ or cell cycle activity of HSCs (Fig. S4 D,E), blocking cADPR-dependent Ca^2+^ release from the endoplasmic reticulum by the 8-Bromo-cADPR antagonist reduced [Ca^2+^]_c_ in HSCs (Fig. 4E) and promoted their cell cycle entry (Fig. 4F). To confirm that CD38 truly regulates [Ca^2+^]_c_ in HSCs we used thapsigargin (TG), which inhibits Ca^2+^ transport from the cytoplasm into the ER. As expected (Fig 4G), [Ca^2+^]_c_ increased significantly faster in CD38^+^ HSCs compared to CD38^-^ cells (Fig. 4H–I) and this difference was abrogated by treatment with 78c, the CD38 inhibitor (Fig. 4J), suggesting that Ca^2+^ release from the ER in CD38^+^ HSCs is mediated by CD38. Together, these data suggest that CD38-dependent cADPR but not ADPR production contributes to high [Ca^2+^]_c_ concentration in dHSCs, which maintains their quiescence.

Human HSCs (hHSC) are CD38^lo/-^ (Fig. S5 A-B) (32), therefore we investigated whether the CD38 ecto-enzymatic activity at the neighboring CD38-positive cells may regulate hHSC quiescence. Indeed, we cultured hHSCs together with CD38^+^ tumor cell line (Fig. S5B) and found that inhibition of CD38 enzymatic activity by 78c inhibitor (Fig. S5C) led to the cell cycle entrance of hHSCs (Fig. 4K). Therefore, CD38 can regulate hHSC quiescence in a paracrine fashion.

### c-Fos regulates quiescence of dHSCs

To clarify how the CD38/cADPR/Ca^2+^ axis regulates HSC dormancy, we performed a bulk transcriptome RNA sequencing of CD38^+^ and CD38^-^ LT-HSCs (LSK CD48^-^ CD150^+^ CD34^-^ CD201^+^) and found that while 205 genes were significantly down-regulated in CD38^+^ LT-HSCs, 225 were up-regulated (Fig. 5A, Supp. Table 4). Gene set enrichment analysis (GSEA) revealed a significant down-regulation of genes related to hematopoietic stem cell differentiation programs, mitochondrial respiratory chain complex assembly, and NADH dehydrogenase complex in CD38^+^ dHSCs compared to CD38^-^ LT-HSCs. Similarly, gene sets controlling the response to calcium ions, extracellular matrix interaction, and TGF-β1 response were up-regulated in CD38^+^ dHSCs (Fig. 5 B, C). HSC-related genes, such as *Hoxb9, H19, VwF, Clu*, and *Sele* (12, 33–36), as well as genes associated with HSC dormancy, namely, *Gprc5c, Meis2* (6), and *Neo1* (37), were up-regulated in CD38^+^ dHSCs (Fig. 5D). We did not find significant differences in *Cdk2, Cdk4, Cdk6*, and *CyclinD1* expression but CD38^+^ dHSCs expressed the cell cycle inhibitors *Cdkn1a* and *Cdkn1c* at higher levels than CD38^-^ LT-HSCs (Fig. 5E). Intriguingly, the transcription factor *Fos*, whose expression was previously correlated to cell cycle activation (38), was one of the most significantly upregulated genes in CD38^+^ dHSCs, and this observation was further corroborated by the fact that CD38^+^ dHSCs displayed higher levels of transcriptionally active phosphorylated c-Fos (at Threonine 232, p-c-Fos) (39) compared to CD38^-^ LT-HSCs (Fig. 5F, Fig. S4F).

**Figure 5.**
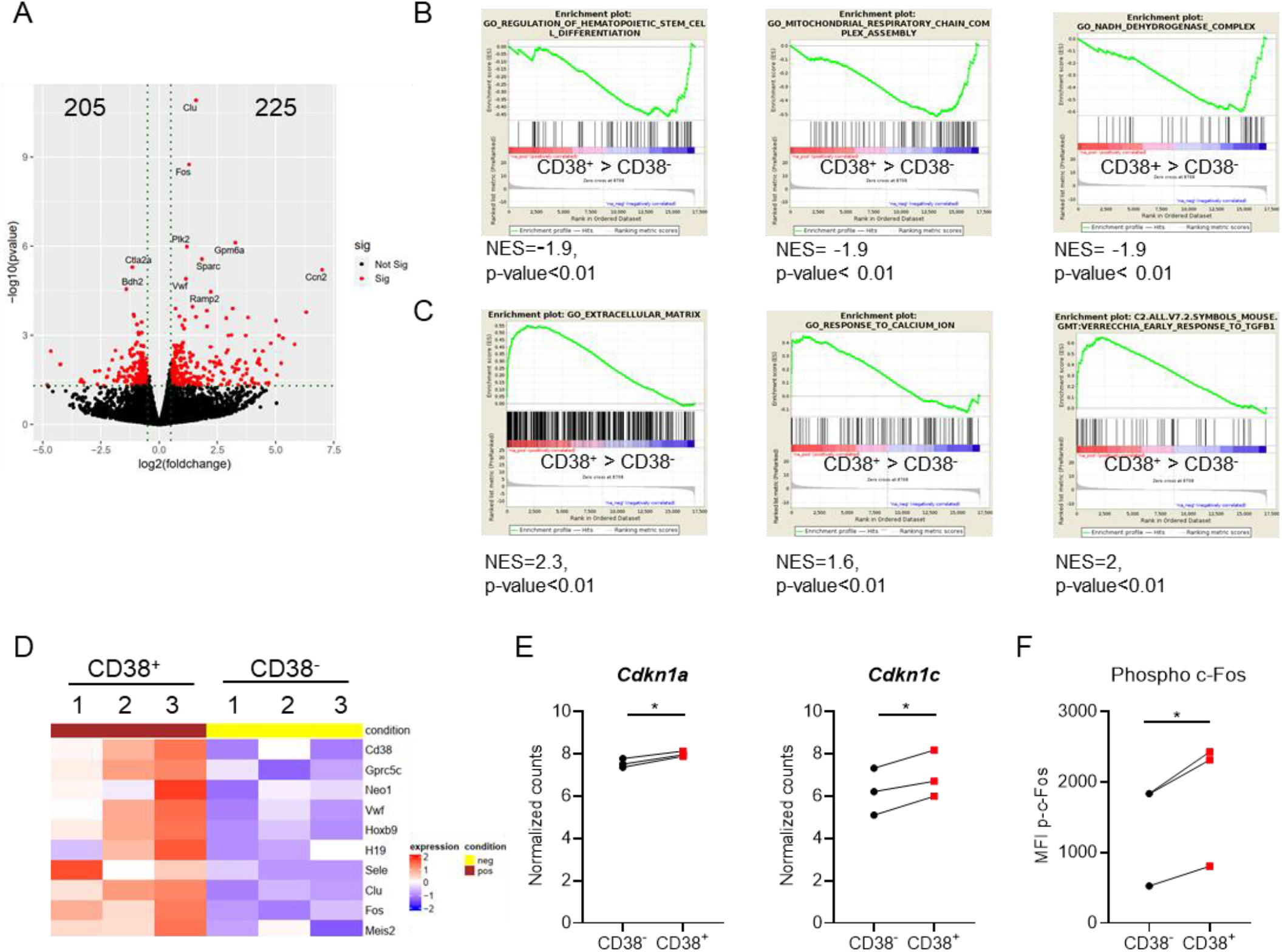
c-Fos is up-regulated in CD38^+^ dHSCs. (A) Volcano plot of differentially expressed genes in CD38^+^ LT-HSCs compared to CD38^-^ cells. (B) GSEA of down-regulated genes in CD38^+^ LT-HSCs compared to CD38^-^ stem cells. (C) GSEA of upregulated genes in CD38^+^ LT-HSCs compared to CD38^-^ cells. (D) Heat map depicting dHSCs and cell cycle-related genes expressed in CD38^+^ and CD38^-^ LT-HSCs. (E) Normalized expression of *Cdkn1a* and *Cdkn1c* in CD38^+^ vs CD38^-^ LT-HSCs. P-values were calculated using the paired *t*-test, *p<0.05. (F) MFI of intracellular p-c-Fos in CD38^+^ and CD38^-^ HSCs. P-values were calculated using the paired *t*-test; *p<0.05.

Correspondingly, single-cell tracking analysis revealed that blocking c-Fos interaction with DNA using a specific inhibitor, T-5224 (40), induced division of CD38^+^ dHSCs but not CD38^-^ LT-HSCs (Fig. 6A–C). Moreover, administering T-5224 to mice led to the partial loss of quiescence in CD38^+^ HSCs but not in CD38^-^ counterparts (Fig. 6D). These data suggest that c-Fos transcriptional activity is necessary for CD38-mediated dHSCs quiescence. As inhibiting the transcriptional activity of c-Fos affected cell cycle entrance of CD38^+^ dHSCs but not CD38^-^ LT-HSCs, we hypothesized that CD38 regulates c-Fos expression via cADPR/Ca^2+^ (Fig. 6E). Indeed, treatment of HSCs with a cADPR antagonist (Br-cADPR) reduced the levels of active p-c-Fos (Fig. 6E), supporting the notion that the CD38/cADPR/Ca^2+^ axis regulates c-Fos levels in dHSCs.

**Figure 6.**
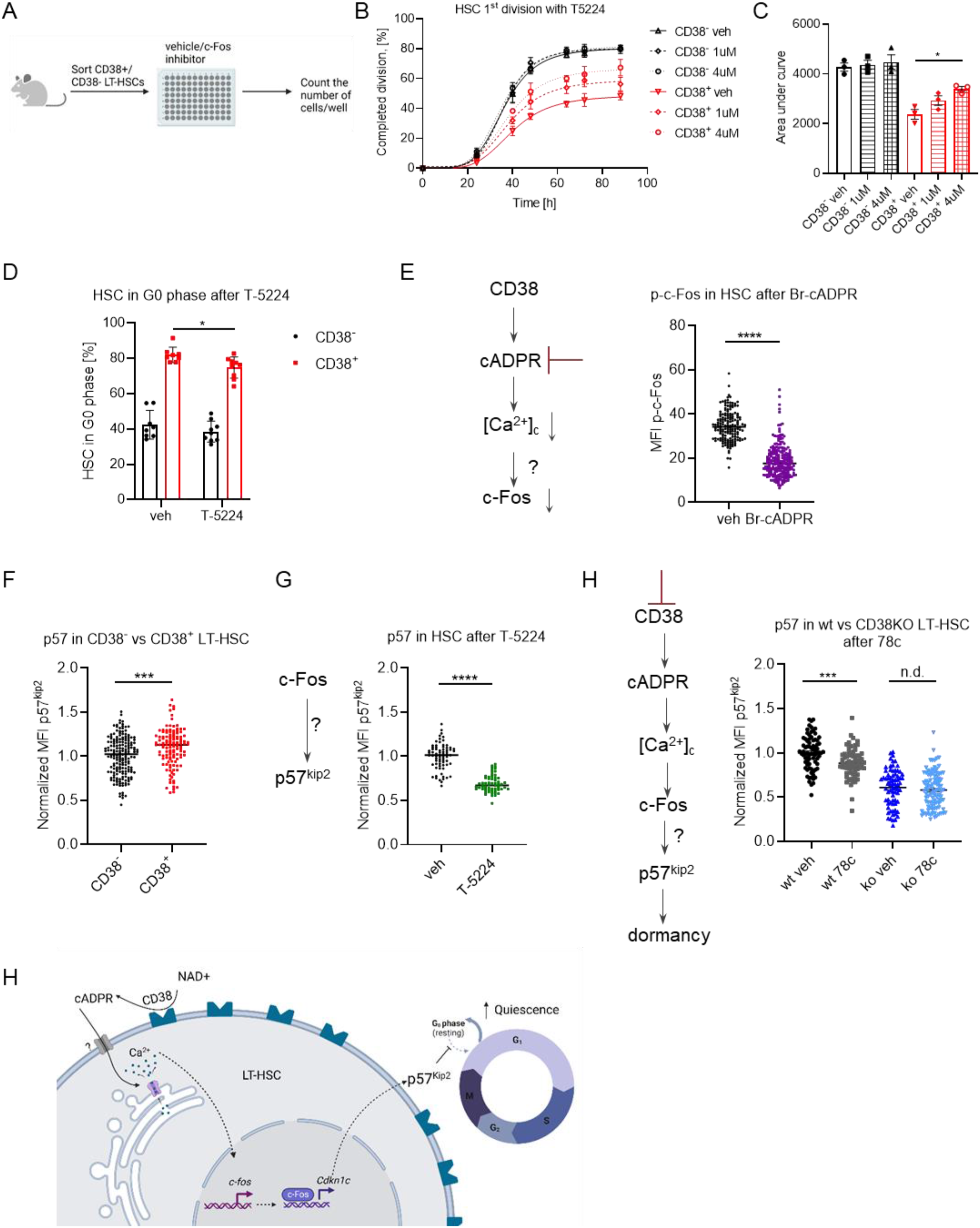
c-Fos maintains dHSC dormancy via p57^kip2^. (A) Single CD38^-^ and CD38^+^ LT-HSCs were sorted and cultured in liquid media with or without the c-Fos inhibitor, T5224. (B) Frequency of LT-HSCs that had completed the first division during incubation is presented (3 independent experiments). (C) Quantification of AUC for data in panel B. P-value was calculated using the paired *t*-test. *p<0.05 (D) Cell cycle analysis of HSCs at 24 h after T-5224 injection, n=8 vs 9, 2 independent experiments. P-value was calculated using the Mann-Whitney *U*-test, *p<0.05 (E) p-c-Fos immunofluorescence in HSCs after 24 h with or without Br-cADPR *in vitro* (n=164 - veh, 219 - Br-cADPR, 2 independent experiments). (F) Quantification of MFI of p57^kip2^ in CD38^+^ dHSCs, and CD38^-^ LT-HSCs (n=181 - CD38^-^, 122 - CD38^+^, 2 independent experiments). (G) p57^kip2^ immunofluorescence in HSCs after 24 h with or without c-Fos inhibitor *in vitro* (n=72 - veh, 61 - T-5224, 2 independent experiments). (H) p57^kip2^ immunofluorescence in CD38+ wt vs CD38KO LT-HSCs after 24 h with or without 78c inhibitor *in vitro* (n=78/68/72/97 for wt veh/wt 78c/ko veh/ko 78c, respectively). For panels E-H, p-values were calculated using unpaired *t*-test. ***p<0.001, ****p<0.0001.

### CD38 controls p57^kip2^ expression in dHSCs via c-Fos

To gain mechanistic insight into how CD38 and c-Fos regulate HSC dormancy, we analyzed the presence of c-Fos binding motifs in the regulatory regions of stem cell-related genes that were upregulated in CD38^+^ dHSCs (Fig. 5D, E) and found that several genes, including *Cdkn1c*, a well-known regulator of HSC quiescence (17), have c-Fos binding motifs (Suppl. Table 5). Therefore, it is possible that c-Fos blocks cell cycle entrance of dHSCs through *Cdkn1c* expression. Indeed, we confirm that not only expression of the *Cdkn1c* gene (Fig. 5E) but also that of its gene product, p57^Kip2^ protein, is higher in CD38^+^ dHSCs than in CD38^-^ LT-HSCs (Fig. 6F). Furthermore, inhibition of the interaction between DNA and c-Fos or the CD38 enzymatic activity led to a reduction in p57^kip2^ protein expression (Fig. 6 G, H), thereby supporting our hypothesis that CD38 activates *Cdkn1c* expression in dHSCs via c-Fos.

## Discussion

We identify CD38 as a novel surface marker that enables isolation of the most dormant population of LT-HSCs. Mechanistically, we demonstrate that CD38 itself regulates stem cell dormancy by shuttling intracellular [Ca^2+^]_c_ in a CD38/cADPR-dependent manner, which results in a consequent increase in c-Fos and the expression of the cell cycle inhibitor p57^kip2^ (Fig. 6H).

The gene expression profile of single HSCs revealed that their heterogeneity is predominantly determined by genes related to the cell cycle, which is in agreement with previous findings (41). We found that *Cd38* expression is inversely correlated with that of cell cycle activators but positively associated with the cell cycle inhibitor *Cdkn1c* and other well-known genes that define the most quiescent LT-HSCs, such as *VwF, Procr, Fgd5*, and *Gprc5c* (6, 12, 13, 15). Importantly, our results indicate that CD38^+^ dHSCs reside at the top of the hematopoietic hierarchy, that they have the highest repopulation capacity upon serial transplantation, and that they are the most quiescent stem cells both under steady-state conditions and in response to a variety of hematopoietic perturbations, i.e., they have the characteristics of dHSCs (4). Hence, CD38 represents a marker that will help circumvent the limitations of the long-term label-retaining assays (4, 5, 24, 42) or even negate the necessity of reporter mice (6, 9) for studying the mechanisms underlying HSC dormancy. Notably, while it is established that the bone marrow niche cells maintain HSCs in the quiescent state (43), little is known about which niche cells are maintaining HSCs dormant. The use of CD38 will allow dHSC imaging to clarify the localization of dHSCs in bone marrow niche. Moreover, CD38 is an ecto-enzyme, and we describe, for the first time, the mechanism involved in HSCs’ dormancy, i.e., autocrine regulation via cADPR. Nonetheless, both substrates and products of its enzymatic activity could also potentially regulate the niche cells neighboring dHSCs, suggesting that dHSCs may also actively modulate their local microenvironment in a paracrine fashion.

Indeed, we found that CD38 enzymatic activity on neighboring cells regulated the proliferation of CD38-negative hHSCs. Interestingly, several hematological malignancies (chronic myeloid leukemia, acute myeloid leukemia, acute lymphoblastic leukemia, and multiple myeloma) express CD38 at a high level (44). We hypothesize that tumour microenvironment enriched in the products of CD38 ecto-enzymatic activity may keep healthy HSCs in the quiescent state leading to cancer-related pancytopenia as well as it may preserve the dormancy of cancer stem cells leading to disease persistence. Therefore, a better understanding of the mechanisms controlling human HSC dormancy is required to support healthy hematopoiesis in patients with hematologic malignancies and develop more powerful strategies for cancer eradication.

Recently, Fukushima et al. have shown that high intracellular [Ca^2+^] prevents cell cycle entry of LT-HSCs (9). In line with this finding, we show that CD38^+^ dHSCs had higher [Ca^2+^]_c_ than CD38^-^ LT-HSCs, that cADPR, the product of CD38/NAD catabolic activity (27), is responsible for the high [Ca^2+^]_c_ in LT-HSCs, and that it directly correlates with the up-regulation of *Fos* gene expression and phosphorylated c-Fos protein. Importantly, these observations are in agreement with the finding that the upstream regulatory region of the *Fos* gene contains a cyclic-AMP response element (CRE) DNA motif, which is critical for the activation of *Fos* transcription by Ca^2+^ (46).

Generally, *Fos* is known as one of the immediate early genes that is transiently expressed in stimulated cells, leads to cell cycle progression (47, 48), and is a positive regulator of myeloid differentiation (49). Besides, c-*Fos* is an oncogene, whose expression is often upregulated in hematologic malignancies, e.g., in chronic and acute myeloid leukemia (50, 51). In contrast, our data clearly reveal that inhibiting the interaction between c-Fos and DNA in dHSCs reduced protein levels of the cell cycle inhibitor p57^kip2^ and stimulated cell cycle entry. Therefore, c-Fos can activate multiple transcriptional programs in a cell type-specific manner and as its role in hematopoiesis regulation is very complex, further investigations are required.

In summary, we reveal that the CD38/cADPR/Ca^2+^/c-Fos/p57^kip2^ axis regulates HSC dormancy. Manipulation of this axis can potentially stimulate dHSC expansion and their efficient response to a variety of hematopoietic stresses. Importantly, the introduction of a new surface marker for the isolation of dHSCs opens up avenues for future research addressing mechanisms that regulate dormant and active states.

## Methods

### Reagents and resources

Supplemental Table 6 lists all reagents used.

### Mice

C57BL/6N (B6), B6.SJL-Ptprc^a^Pep3^b^/BoyJ (SJL) were purchased from The Jackson Laboratory. C57BL/6N (B6) and B6.SJL-Ptprc^a^Pep3^b^/BoyJ (SJL) mice were crossed to produce F1 progeny (CD45.1/CD45.2) for transplantation experiments. To study the division history of HSCs, R26^rtTA^/*Col1A1*^H2B-GFP^ mice were used (5); the induction of H2B-GFP expression was performed as described in (24). B6.129P2-Cd38^tm1Lnd^/J (CD38KO) mice were obtained from Dr. Jaime Sancho and Dr. Frances Lund. Ki67^RFP^ knock-in mice have been described previously (52, 53). All mice were bred and maintained under specific pathogen–free conditions in the animal facility at the Medical Theoretical Center of the Technical University, Dresden, Germany. Experiments were performed in accordance with the German animal welfare legislation and were approved by the “Landesdirektion Sachsen”. Unless specified otherwise, 8–16 week-old mice of both genders were used for experiments.

### Cell isolation and flow cytometry

Cells were isolated from tibiae, femora, pelvis, and vertebrae by crushing bones in 5% fetal bovine serum (FBS) in phosphate-buffered saline (PBS) and passing them through a 40-um filter. Erythrocytes were lysed using ACK Lysing Buffer. To calculate the amount of cells, cells isolated from 2 tibiae and 2 femora were stained with DAPI at 0.1 ug/ml and DAPI-negative cells were counted using MACSQuant Analyzer (Myltenyi Biotec). For sort, cells were stained with c-Kit bio antibody and Anti-Biotin MicroBeads were added to enrich for c-Kit^+^ cells using LS columns. HSCs were defined as Lin^-^ (negative for B220, CD3ε, CD19, NK1.1, Gr1, Ter119, and CD11b) Sca1^+^ c-Kit^+^ (LSK) CD48^-^ CD150^+^ cells. LT-HSCs: LSK CD48^-^ CD150^+^ CD34^-^ CD201^+^, MPP2: LSK CD48^+^ CD150^+^, MPP3/4: LSK CD48^+^ CD150^-^, and ST-HSCs: LSK CD48^-^ CD150^-^. Granulocyte-monocyte progenitors (GMP) were defined as Lin^-^ Sca1^-^ c-Kit^+^ (LK) CD16/32^+^ CD150^-^, pre-megakaryocyte progenitors, PreMeg: LK CD16/32^-^ CD41^+^ CD150^+^, colony forming unit-erythroid, CfuE: LK CD16/32^-^ CD41^-^ CD105^+^ CD150^-^, pre-colony forming unit-erythroid, PreCfuE: LK CD16/32^-^ CD41^-^ CD105^+^ CD150^+^, megakaryocyte-erythroid progenitors, MEP: LK CD16/32^-^ CD41^-^ CD105^-^ CD150^+^, common myeloid progenitors, CMP: LK CD16/32^-^ CD41^-^ CD105^-^ CD150^-^. All analyses were performed on a BD LSR II, BD FACSAria II, BD LSRFortessa X-20, or a BD FACSCanto II (BD Bioscience). Data were analyzed using FlowJo software 10.7.1 (BD Bioscience).

### Single cell RNA sequencing with 10x Genomics and analysis

LT-HSCs (LSK CD48^-^ CD150^+^, 3000 cells) from four B6 mice were sorted into BSA-coated tube containing 5 μl of PBS with 0.04% BSA. All cells were carefully mixed with reverse transcription mix before loading them in a Chromium Single Cell A Chip on the 10X Genomics Chromium system (54) and processed further following the guidelines of the 10x Genomics user manual for Chromium Single Cell 3’ RNA-seq Library v2. In short, the droplets were directly subjected to reverse transcription, the emulsion was broken and cDNA was purified using silane beads. After amplification of cDNA with 12 cycles, it underwent a purification with 0.6 volume of SPRI select beads. After quality check and quantification, 20 μl cDNA was used to prepare single cell RNA-seq libraries - involving fragmentation, dA-Tailing, adapter ligation and a 13 cycles indexing PCR based on manufacturer’s guidelines. After quantification, both libraries were sequenced on an Illumina Nextseq500 system in paired-end mode with 26 bp/57 bp (for read 1 and 2 respectively), thus generating ~50-80 mio. fragments for the transcriptome library on average. The raw count data generated from Cell Ranger pipeline was processed using Seurat v3.1 (10) by following the standard pipeline. Cells were filtered based on quality metrics (number of genes, total UMI counts, percentage of mitochondrial genes). Subsequently, filtered data were merged using “FindIntegrationAnchors” function of Seurat 3. For further analysis, merged data were log-normalized, regressed for library size, and percentage of mitochondrial genes and scaled. Cell cycle and dormancy scores were calculated with G2M and S phase genes from Seurat package and dormancy related genes (6) (see Suppl. Table 1) and using “AddModuleScore” function of Seurat 3. For pseudotime trajectory analysis, standard pipeline of Monocle 2 (54) was used and dimensionality reduction was performed using “reduceDimension” function of Monocle 2 with following parameters: num_dim = 10, norm_method = “log”, reduction_method = “tSNE”, residualModelFormulaStr = “~Age”. To visualize gene modules showing similar kinetic trends, “plot_pseudotime_heatmap” function was used accounting a list of genes showing significant score for differential expression along pseudotime (q-value < 0.05) and genes that change as a function of pseudotime were grouped in three clusters.

### Gene ontology analysis

GO term analysis was performed using Enrichr (55). Complete gene list per clusters were obtained by using the “crisp gene set”. For visualization, statistically significant (p-value < 0.05) terms were selected from top five pathways (see Supp. Table 3).

### LT-HSCs transplantation

For primary transplantation, 50 CD38^-^ or CD38^+^ LT-HSCs (LSK CD48^-^ CD150^+^ CD34^-^ CD201^+^) were sorted and transplanted together with 5×10^5^ total BM competitor cells. For secondary transplantation, 5×10^6^ CD45^+^ total BM cells were transplanted into lethally irradiated recipients. Recipient mice were lethally irradiated (9 Gy), and the cells were injected i.v..

### Cell cycle analyses

Cell cycle was analyzed using staining for Ki67 and DAPI as described before (26). To label dHSCs, mice were injected with 1 mg of BrdU i.p. and kept with 0.8 mg BrdU per 1 ml in drinking water for 14 more days before sacrificing. Water was changed every 3 days. BrdU incorporation analysis was performed using anti-BrdU antibody as described before (19).

### *In vivo* stress models and drug administration

To mimic viral infection, pIC was administered i.p. at a dose of 5 mg/kg 18 or 48 h prior to analysis. To selectively deplete platelets, rabbit anti-mouse antiplatelet serum was injected i.p. 18 h prior to analysis. The effective dose of anti-platelet serum was determined before the use in experiments (26), and a dose resulting in <150×10^3^ platelets per microliter of blood at 2 h after the injection was considered suitable for experiments. Control mice were injected with the corresponding amount of normal rabbit serum. To study HSC response to chemotherapy, 5-FU was injected i.p. at a dose of 150 mg/kg 4 or 8 days prior to analysis. To study the effects of c-Fos inhibition, T-5224 in the vehicle (2% DMSO+30% PEG300+2% Tween80 in ddH2O) was injected i.p. at 30 mg/kg/mouse 18 h prior to analysis. Control mice were injected with the corresponding amount of vehicle.

### Intracellular calcium staining and flux

Calcium staining and flux were estimated by flow cytometry. Cells were sequentially incubated with 0.3 μM Fluo-8 AM for 1 h at room temperature and with 2 μM Indo-1 and 0.02% Pluronic F-127 in HBSS for 30 min at 37°C, washed, and resuspended in HBSS. Wavelength filters for 405±20nm (violet emission) and 530±30nm (450 LP filter, blue emission) were used to visualize Ca^2+^-bound and -unbound dye ratio by flow cytometry, respectively. After recording baseline calcium, thapsigargin (TG, 1 mM) was added to the sample to induce Ca^2+^ flux. Alternatively, 78c (1 μM) was added to cells 5 min before TG. The average ratio, R, of bound/unbound Indo-1 (405nm/485nm emission) was calculated.

### Single cell LT-HSCs *in vitro* culture

Single long term-HSCs were sorted into 96-well plates containing StemSpan medium with 10 ng/mL SCF, 10 ng/mL THPO, 20 ng/mL IGF2, and 10 ng/mL FGF1, with or without 1 μM / 4 μM c-Fos inhibitor T-5224, 0.1 μM CD38 inhibitor 78c, and cultured for 3 days at 37°C with 5% CO2. The number of the cells per well was monitored twice daily under a light microscope.

### *In vitro* treatment of LSK cells

LSK cells from Ki67^RFP^ mice were sorted and cultured (5 × 10^4^ per well) in StemSpan medium with 10 ng/mL SCF, 10 ng/mL THPO, 20 ng/mL IGF2, and 10 ng/mL FGF1 with or without 25 or 100 uM 8-Br-cADPR or 25-100 uM 8-Br-ADPR or 4 uM T5224. 24 h later cells were stained with anti CD48, CD150, Kit, Sca-1 antibodies, and 0.3 uM Fluo-8 AM. Alternatively, surface stained cells were fixed using eBioscience kit and HSCs were sorted on glass slides for immunofluorescent staining.

### Immunofluorescent staining

HSCs sorted on glass slides were used for the immunofluorescent staining. Cells were blocked with 20% horse serum in 1X Permeabilization buffer (eBioscience) for 30 min at RT, stained with rabbit anti-phospho-c-Fos or rabbit anti-p57 antibodies for 2h, washed and then incubated with secondary anti-rabbit AlexaFluor 488 antibody for 30 min. Cells were mounted using DAPI-containing mounting and sealed. Images were captured using a Leica TCS SP5 confocal microscope (Leica Microsystems) using 63x objective. From 6 to 8 z-stacks were taken per image and fluorescence was analyzed using Fiji (56).

### Human HSC *in vitro* culture

Bone marrow stem cell apheresates were collected from healthy donors at the Dresden Bone Marrow Transplantation Centre of the University Hospital Carl Gustav Carus. The donors fulfilled the standards for bone marrow donation (e.g. free of HIV, HBV, and serious illness), were informed and gave their approval. The study was approved by the local ethics commission (EK263122004, EK114042009). MOLM-13 were obtained from ATCC. Mononuclear cells (MNCs) were isolated using Ficoll fractioning. Briefly, BM aspirates were layered on top of Ficoll-Paque PLUS media and centrifuged at 800g for 20 min at 20°C (brake off), MNCs were then isolated from buffy coat fraction in the interphase of Ficoll gradient. MNCs were incubated with anti-CD34 MicroBeads and enriched for CD34+ cells using LS columns. MNCs were stained for surface CD34 and CD38, and 3000 HSCs (CD38^lo/-^ CD34+) were sorted into 96-well plates into CellGenix TM GMP SCGM medium supplemented with 2.5% FBS, human FLT3L, human SCF, and human IL-3 (all 2.5 ng/ml) containing 1×10^5^ MOLM-13 cells, and cultivated at 37°C 5% CO2 in duplicates with or without 1 uM 78c. Next day, the cells were stained with anti CD34, CD38 antibodies. Cell cycle was analyzed using staining for Ki67 and DAPI as described before (26).

### Bulk RNA sequencing

A total of 2000 LT-HSCs (LSK CD48^-^ CD150^+^ CD34^-^ CD201^+^) that were CD38^+^ or CD38^-^ were sorted (pooled cells from 10 mice per sample). Bulk RNA sequencing was performed as previously described (26). Illumina sequencing was done on a Nextseq500 with an average sample sequencing depth of 60 million reads.

### Transcriptome Mapping

Low quality nucleotides were removed with Illumina fastq filter (http://cancan.cshl.edu/labmembers/gordon/fastq_illumina_filter/). Reads were further subjected adaptor trimming using cutadapt (57). Alignment of the reads to the Mouse genome was done using STAR Aligner (58) using the parameters: “--runMode alignReads -- outSAMstrandField intronMotif --outSAMtype BAM SortedByCoordinate --readFilesCommand zcat”. Mouse Genome version GRCm38 (release M12 GENCODE) was used for the alignment. HTSeq-0.6.1p1 (59) was used to count the reads that map to the genes in the aligned sample files. Read Quantification was performed using the parameters: ‘htseq-count -f bam -s reverse -m union -a 20’. The GTF file (gencode.vM12.annotation.gtf) used for read quantification was downloaded from Gencode (60).

### Differential Expression Analysis

Gene centric differential expression analysis was performed using DESeq2_1.8.1 (61). Volcano plot was created using ggplot2_1.0.1 (62). Heatmaps were generated using ComplexHeatmap package of R/Bioconductor (63).

### Gene enrichment analysis

Pathway and functional analyses were performed using GSEA (64). GSEA is a stand-alone software with a GUI. To run GSEA, a ranked list of all the genes from DESeq2 based calculations was created by taking the -log10 of the p-value and multiplying it with the sign the of the fold change. This ranked list was then queried against MsigDB (65).

### Data

For original data, please contact the corresponding author. Bulk and single-cell RNA-sequencing data are available at GEO under accession numbers GSE196760 and GSE196759 respectively.

### Statistics

Data are presented as mean ± SEM. Significance was calculated using the Mann-Whitney U test, unless stated otherwise. All statistical analyses were performed using GraphPad Prism 8.2.1 for Windows (GraphPad Software, La Jolla, CA; www.graphpad.com).

## Study approval

Animal experiments were performed in accordance with the German animal welfare legislation and were approved by the “Landesdirektion Sachsen”.

Additional and detailed descriptions of procedures can be found in supplemental Methods.

## Author contributions

T.G. designed the study and supervised research; L.I., T.G., performed most of the experiments, analyzed, and interpreted data and wrote the manuscript; A.G. performed the long-term label retention experiment; J.A.P.V., L.M. and R.W. performed experiments; S.P.S., S.E.E. analyzed and interpreted single-cell RNA sequencing data; A.S. analyzed bulk deep sequencing data; M.W. and M. von B. participated in scientific discussion and data interpretation; S.R and A.D. performed next generation sequencing; J.S. and F.L. provided CD38KO mice; B.W. contributed to the study design and edited the manuscript; T.G., B.W., M.B., and M.S. organized research and interpreted data. T.C. interpreted data and edited the manuscript. All authors discussed the results and commented on the manuscript.

## Acknowledgements

The figures 2D, 3C, 3D, 4B, 4G, 4K, 6A, 6H, S3G, S3L were created using BioRender.com. We thank Anja Krüger and Robert Kuhnert for technical assistance, Core Facility Cellular Imaging at Faculty of Medicine, TU Dresden as well as Deep Sequencing Facility, DRESDEN-*concept* Genome Center. This work was supported by a grant from the DFG (GR 4857/2-1) and Mildred-Scheel-Nachwuchszentrum fellowship to T.G. and WI3291/5-1, 12-1 and 13-1 to B.W. T.C. is supported by the DFG (TRR332, project B4). S.P.S. was supported by FNRS (40005588 – MISU-PROL) and Jaumotte-Demoulin Foundation. S.E.E. was supported by PhD Fellowship from FNRS (40006730 – ASP). English language and content editing was provided by Vasuprada Iyengar.

## References

1. Glimm H, et al. Human hematopoietic stem cells stimulated to proliferate in vitro lose engraftment potential during their S/G(2)/M transit and do not reenter G(0). Blood. 2000;96(13):4185–93.

2. Nygren JM, and Bryder D. A novel assay to trace proliferation history in vivo reveals that enhanced divisional kinetics accompany loss of hematopoietic stem cell self-renewal. PLoS One. 2008;3(11):e3710.

3. Qiu J, et al. Divisional history and hematopoietic stem cell function during homeostasis. Stem Cell Reports. 2014;2(4):473–90.

4. Wilson A, et al. Hematopoietic stem cells reversibly switch from dormancy to self-renewal during homeostasis and repair. Cell. 2008;135(6):1118–29.

5. Foudi A, et al. Analysis of histone 2B-GFP retention reveals slowly cycling hematopoietic stem cells. Nat Biotechnol. 2009;27(1):84–90.

6. Cabezas-Wallscheid N, et al. Vitamin A-Retinoic Acid Signaling Regulates Hematopoietic Stem Cell Dormancy. Cell. 2017;169(5):807–23 e19.

7. Essers MA, et al. IFNalpha activates dormant haematopoietic stem cells in vivo. Nature. 2009;458(7240):904–8.

8. Zhang YW, et al. Hyaluronic acid–GPRC5C signalling promotes dormancy in haematopoietic stem cells. Nature Cell Biology. 2022;24(7):1038–48.

9. Fukushima T, et al. Discrimination of Dormant and Active Hematopoietic Stem Cells by G0 Marker Reveals Dormancy Regulation by Cytoplasmic Calcium. Cell Rep. 2019;29(12):4144–58 e7.

10. Stuart T, et al. Comprehensive Integration of Single-Cell Data. Cell. 2019;177(7):1888–902 e21.

11. Hao Y, et al. Integrated analysis of multimodal single-cell data. Cell. 2021;184(13):3573–87 e29.

12. Sanjuan-Pla A, et al. Platelet-biased stem cells reside at the apex of the haematopoietic stem-cell hierarchy. Nature. 2013;502(7470):232–6.

13. Balazs AB, et al. Endothelial protein C receptor (CD201) explicitly identifies hematopoietic stem cells in murine bone marrow. Blood. 2006;107(6):2317–21.

14. Wilson NK, et al. Combined Single-Cell Functional and Gene Expression Analysis Resolves Heterogeneity within Stem Cell Populations. Cell Stem Cell. 2015;16(6):712–24.

15. Gazit R, et al. Fgd5 identifies hematopoietic stem cells in the murine bone marrow. J Exp Med. 2014;211(7):1315–31.

16. Zou P, et al. p57(Kip2) and p27(Kip1) cooperate to maintain hematopoietic stem cell quiescence through interactions with Hsc70. Cell Stem Cell. 2011;9(3):247–61.

17. Matsumoto A, et al. p57 is required for quiescence and maintenance of adult hematopoietic stem cells. Cell Stem Cell. 2011;9(3):262–71.

18. Yamazaki S, et al. TGF-beta as a candidate bone marrow niche signal to induce hematopoietic stem cell hibernation. Blood. 2009;113(6):1250–6.

19. Grinenko T, et al. Clonal expansion capacity defines two consecutive developmental stages of long-term hematopoietic stem cells. J Exp Med. 2014;211(2):209–15.

20. Morita Y, et al. Heterogeneity and hierarchy within the most primitive hematopoietic stem cell compartment. J Exp Med. 2010;207(6):1173–82.

21. Morcos MNF, et al. SCA-1 Expression Level Identifies Quiescent Hematopoietic Stem and Progenitor Cells. Stem Cell Reports. 2017;8(6):1472–8.

22. Rabe JL, et al. CD34 and EPCR coordinately enrich functional murine hematopoietic stem cells under normal and inflammatory conditions. Exp Hematol. 2020;81:1–15 e6.

23. Oguro H, et al. SLAM family markers resolve functionally distinct subpopulations of hematopoietic stem cells and multipotent progenitors. Cell Stem Cell. 2013;13(1):102–16.

24. Morcos MNF, et al. Continuous mitotic activity of primitive hematopoietic stem cells in adult mice. J Exp Med. 2020;217(6).

25. Liang R, et al. Restraining Lysosomal Activity Preserves Hematopoietic Stem Cell Quiescence and Potency. Cell Stem Cell. 2020;26(3):359–76 e7.

26. Ramasz B, et al. Hematopoietic stem cell response to acute thrombocytopenia requires signaling through distinct receptor tyrosine kinases. Blood. 2019;134(13):1046–58.

27. Graeff R, et al. Mechanism of cyclizing NAD to cyclic ADP-ribose by ADP-ribosyl cyclase and CD38. J Biol Chem. 2009;284(40):27629–36.

28. Tarrago MG, et al. A Potent and Specific CD38 Inhibitor Ameliorates Age-Related Metabolic Dysfunction by Reversing Tissue NAD(+) Decline. Cell Metab. 2018;27(5):1081–95 e10.

29. Huang C, et al. Extracellular Adenosine Diphosphate Ribose Mobilizes Intracellular Ca2+ via Purinergic-Dependent Ca2+ Pathways in Rat Pulmonary Artery Smooth Muscle Cells. Cellular Physiology and Biochemistry. 2015;37(5):2043–59.

30. Ernst IM, et al. Adenine Dinucleotide Second Messengers and T-lymphocyte Calcium Signaling. Front Immunol. 2013;4:259.

31. Lee HC. Cyclic ADP-ribose: a new member of a super family of signalling cyclic nucleotides. Cell Signal. 1994;6(6):591–600.

32. Reya T, et al. Stem cells, cancer, and cancer stem cells. Nature. 2001;414(6859):105–11.

33. Copley MR, et al. The Lin28b-let-7-Hmga2 axis determines the higher self-renewal potential of fetal haematopoietic stem cells. Nat Cell Biol. 2013;15(8):916–25.

34. Kumar P, et al. HMGA2 promotes long-term engraftment and myeloerythroid differentiation of human hematopoietic stem and progenitor cells. Blood Adv. 2019;3(4):681–91.

35. Pineault N, et al. Differential expression of Hox, Meis1, and Pbx1 genes in primitive cells throughout murine hematopoietic ontogeny. Exp Hematol. 2002;30(1):49–57.

36. Venkatraman A, et al. Maternal imprinting at the H19-Igf2 locus maintains adult haematopoietic stem cell quiescence. Nature. 2013;500(7462):345–9.

37. Renders S, et al. Niche derived netrin-1 regulates hematopoietic stem cell dormancy via its receptor neogenin-1. Nat Commun. 2021;12(1):608.

38. Brown JR, et al. Fos family members induce cell cycle entry by activating cyclin D1. Mol Cell Biol. 1998;18(9):5609–19.

39. Monje P, et al. Regulation of the transcriptional activity of c-Fos by ERK. A novel role for the prolyl isomerase PIN1. J Biol Chem. 2005;280(42):35081–4.

40. Aikawa Y, et al. Treatment of arthritis with a selective inhibitor of c-Fos/activator protein-1. Nat Biotechnol. 2008;26(7):817–23.

41. Kowalczyk MS, et al. Single-cell RNA-seq reveals changes in cell cycle and differentiation programs upon aging of hematopoietic stem cells. Genome Res. 2015;25(12):1860–72.

42. Challen GA, and Goodell MA. Promiscuous expression of H2B-GFP transgene in hematopoietic stem cells. PLoS One. 2008;3(6):e2357.

43. Pinho S, and Frenette PS. Haematopoietic stem cell activity and interactions with the niche. Nature Reviews Molecular Cell Biology. 2019;20(5):303–20.

44. Konen JM, et al. The Good, the Bad and the Unknown of CD38 in the Metabolic Microenvironment and Immune Cell Functionality of Solid Tumors. Cells. 2019;9(1):52.

45. Miraki-Moud F, et al. Acute myeloid leukemia does not deplete normal hematopoietic stem cells but induces cytopenias by impeding their differentiation. Proceedings of the National Academy of Sciences. 2013;110(33):13576–81.

46. J.Y.H C, et al. Phosphorylation of transcription factor CREB mediates c-fos induction elicited by sustained hypertension in rat nucleus tractus solitarii. Neuroscience. 1999;88(4):1199–212.

47. Kovary K, and Bravo R. The jun and fos protein families are both required for cell cycle progression in fibroblasts. Mol Cell Biol. 1991;11(9):4466–72.

48. Miao GG, and Curran T. Cell transformation by c-fos requires an extended period of expression and is independent of the cell cycle. Mol Cell Biol. 1994;14(6):4295–310.

49. Lord KA, et al. Proto-oncogenes of the fos/jun family of transcription factors are positive regulators of myeloid differentiation. Mol Cell Biol. 1993;13(2):841–51.

50. Kesarwani M, et al. Targeting c-FOS and DUSP1 abrogates intrinsic resistance to tyrosine-kinase inhibitor therapy in BCR-ABL-induced leukemia. Nat Med. 2017;23(4):472–82.

51. Velten L, et al. Identification of leukemic and pre-leukemic stem cells by clonal tracking from single-cell transcriptomics. Nature Communications. 2021;12(1).

52. Grinenko T, et al. Hematopoietic stem cells can differentiate into restricted myeloid progenitors before cell division in mice. Nat Commun. 2018;9(1):1898.

53. Basak O, et al. Mapping early fate determination in L gr5 + crypt stem cells using a novel K i67-RFP allele. The EMBO Journal. 2014;33(18):2057–68.

54. Qiu X, et al. Single-cell mRNA quantification and differential analysis with Census. Nat Methods. 2017;14(3):309–15.

55. Chen EY, et al. Enrichr: interactive and collaborative HTML5 gene list enrichment analysis tool. BMC Bioinformatics. 2013;14:128.

56. Schindelin J, et al. Fiji: an open-source platform for biological-image analysis. Nat Methods. 2012;9(7):676–82.

57. Martin M. Cutadapt removes adapter sequences from high-throughput sequencing reads. EMBnetjournal. 2011;17(1):10.

58. Dobin A, et al. STAR: ultrafast universal RNA-seq aligner. Bioinformatics. 2013;29(1):15–21.

59. Anders S, et al. HTSeq--a Python framework to work with high-throughput sequencing data. Bioinformatics. 2015;31(2):166–9.

60. GENCODE. GENCODE reference annotation for the human and mouse genomes, Release M12 (GRCm38.p5). https://www.gencodegenes.org/mouse/release_M12.html.

61. Anders S, and Huber W. Differential expression analysis for sequence count data. Genome Biol. 2010;11(10):R106.

62. Wickham H. ggplot2: elegant graphics for data analysis. Springer-Verlag New York; 2016.

63. Gu Z, et al. Complex heatmaps reveal patterns and correlations in multidimensional genomic data. Bioinformatics. 2016;32(18):2847–9.

64. Subramanian A, et al. Gene set enrichment analysis: a knowledge-based approach for interpreting genome-wide expression profiles. Proc Natl Acad Sci U S A. 2005;102(43):15545–50.

65. Liberzon A, et al. Molecular signatures database (MSigDB) 3.0. Bioinformatics. 2011;27(12):1739–40.

