## Supplementary information for "CD38 promotes hematopoietic stem cell dormancy via c-Fos"

**Supplementary information** Ibneeva et al.

### Supplemental Methods

#### Mitochondrial membrane potential and mass

Cells were subjected to 250 nM Tetramethylrhodamine ethyl ester (TMRE) to measure mitochondrial membrane potential (MMP) or to 100 nM Mitotracker Green to measure mitochondrial mass in the presence of 50 µM Verapamil at 37˚C for 20 minutes. Cells were washed and analyzed by flow cytometry.

#### Blood counts

A Sysmex XT-3000 Vet automated hematology analyzer was used to measure blood cell counts.

#### CD38KO transplantation

For LT-HSCs primary transplantation, 50 wt or CD38KO LT-HSCs (LSK CD48$^-$ CD150$^+$ CD34$^-$ CD201$^+$) were sorted and transplanted together with $5\times10^5$ total BM competitor cells. For secondary transplantation, donor HSCs (LSK CD48$^-$ CD150$^+$) were sorted and transplanted together with $5\times10^5$ total BM competitor cells into lethally irradiated recipients. For TBM primary transplantation, $1\times10^6$ CD45+ total BM from wt or CD38KO mice was transplanted together with $1\times10^6$ total BM competitor cells. For secondary transplantation, $5\times10^6$ CD45$^+$ total BM cells were transplanted into lethally irradiated recipients.

#### Transcription Factor Binding Site prediction

"runFimo" command of "memes" https://bioconductor.org/packages/memes package (from R) was run of the 3Kb upstream regions of the genes using the JASPAR2020 database (1).

#### CD38 cyclase activity

CD38 cyclase activity was measured according to (2) with minor changes. MOLM-13 cells were sonicated with 40W 20 kHz 3 times for 5 sec on ice in a lysis buffer containing 10 mM Tris/HCl, pH 7.4, 0.25 M sucrose, 20 mM NaF, 1 mM DTT, 5 mM EDTA, 1 mM PMSF and a protease inhibitor cocktail (chymostatin, leupeptin, antipain and pepstatin A; all chemicals from Sigma–Aldrich). Fluorescence (ex=300 nm, em=410 nm) of reaction mix containing $1\text{x}10^6$ lysed MOLM-13, 200 µM nicotinamide guanine dinucleotide sodium salt (NGD) and 0 - 1.28 µM 78c was recorded for 1 h. To generate dose-response curve, normalized relative fluorescence intensity at 1 h was plotted against log 78c concentration.

**Suppl. Table 5. c-Fos binding motifs in up-regulated stem cell-related genes in CD38$^+$ dHSCs.**

| Name | Start | End | P-value | Matched sequence |
|---|---|---|---|---|
| H19 | 534 | 541 | 6.10E-05 | ATGAGTCA |
| Clu | 142 | 149 | 3.05E-05 | GTGACTCA |
| Cd38 | 7 | 14 | 1.53E-05 | GTGAGTCA |
| Cd38 | 1086 | 1093 | 6.10E-05 | ATGAGTCA |
| Gprc5c | 2338 | 2345 | 7.63E-05 | GTGATTCA |
| Cdkn1c | 522 | 529 | 3.05E-05 | GTGACTCA |
| Cdkn1c | 1999 | 2006 | 3.05E-05 | GTGACTCA |
| Sele | 2928 | 2935 | 1.53E-05 | GTGAGTCA |
| Neo1 | 598 | 605 | 4.58E-05 | GTGAGTAA |
| Neo1 | 2230 | 2237 | 3.05E-05 | GTGACTCA |

**Suppl. Table 6. Reagents and resources**

| Antibodies for flow cytometry and immunofluorescent staining | | | | |
|---|---|---|---|---|
| **Antibody** | **Host** | **Supplier** | **Clone** | **Cat. number** |
| CD45R (B220) PE-Cyanine7 | rat | Thermo Fisher Scientific | RA3-6B2 | 25-0452-82 |
| CD45R (B220) biotin | rat | Thermo Fisher Scientific | RA3-6B2 | 13-0452-82 |
| CD3e APC | rat | Thermo Fisher Scientific | 145-2C11 | 17-0031-82 |
| CD3e biotin | rat | Thermo Fisher Scientific | 145-2C11 | 13-0031-82 |
| CD11b PE | rat | Thermo Fisher Scientific | M1/70 | 12-0112-82 |
| CD11b biotin | rat | Thermo Fisher Scientific | M1/70 | 13-0118-82 |
| CD16/32 A700 | rat | Thermo Fisher Scientific | 93 | 56-0161-82 |

| | | | | |
|---|---|---|---|---|
| CD19 biotin | rat | Thermo Fisher Scientific | 1D3 | 13-0193-82 |
| CD34 FITC | rat | Thermo Fisher Scientific | RAM34 | 11-0341-85 |
| CD38 FITC | rat | Thermo Fisher Scientific | 90 | 11-0381-82 |
| CD38 PE | rat | Biolegend | 90 | 102708 |
| CD38 PerCP-eFluor 710 | rat | Thermo Fisher Scientific | 90 | 46-0381-82 |
| CD41 PerCP eFluor 710 | rat | Thermo Fisher Scientific | MWReg30 | 46-0411-82 |
| CD45 PE | rat | Thermo Fisher Scientific | 3D-F11 | 12-0451-82 |
| CD45.1 PE-Cyanine5 | rat | Thermo Fisher Scientific | A20 | 15-0453-82 |
| CD45.2 APC-eFluor 780 | rat | Thermo Fisher Scientific | 104 | 47-0454-82 |
| CD48 AlexaFluor 700 | rat | Biolegend | HM48-1 | 103426 |
| CD48 APC | rat | BioLegend | HM48-1 | 103412 |
| CD105 APC | rat | BioLegend | 120414 | MJ7/18 |
| CD150 PE-Cyanine7 | rat | BioLegend | TC15-12F12.2 | 115914 |
| CD117 (c-Kit) APC-eFluor 780 | rat | Thermo Fisher Scientific | 2B8 | 47-1171-82 |
| CD117 (c-Kit) biotin | rat | Thermo Fisher Scientific | 2B8 | 13-1171-85 |
| CD229 PE | rat | Biolegend | lyab3 | 122905 |
| Ly-6G/Ly-6C biotin | rat | Thermo Fisher Scientific | RB6-8C5 | 13-5931-82 |
| Ly-6G/Ly-6C eFluor 450 | rat | Thermo Fisher Scientific | RB6-8C5 | 48-5931-82 |
| Ki67 FITC | rat | Thermo Fisher Scientific | SolA15 | 11-5698-82 |
| NK1.1 biotin | rat | Thermo Fisher Scientific | PK136 | 13-5941-82 |
| Streptavidin eFluor 450 | rat | Thermo Fisher Scientific | n/a | 48-4317-82 |
| Streptavidin APC-eFluor 780 | rat | Thermo Fisher Scientific | n/a | 47-4317-82 |

| | | | | |
|---|---|---|---|---|
| Ly-6A/E (Sca-1) PE-Cyanine5 | rat | Thermo Fisher Scientific | D7 | 15-5981-82 |
| Ter-119 biotin | rat | Thermo Fisher Scientific | TER119 | MA5-17819 |
| phospho-c-Fos (Thr232) | rabbit | Bioss antibodies | polyclonal | bs-3153R |
| p57 Kip2 | rabbit | Abcam | EP2515Y | ab75974 |
| Anti-rabbit IgG Brilliant Violet 421 | donkey | Biolegend | polyclonal | 406410 |
| Anti-rabbit IgG (H+L), F(ab')$_2$ Alexa Fluor 488 | goat | Cell signaling | polyclonal | 4412 |
| CD38 Alexa Fluor 700 | human | Biolegend | HB-7 | 356624 |
| CD34 PE-Cyanine7 | human | Biolegend | 561 | 343616 |

| Reagents | | |
|---|---|---|
| **Reagent** | **Supplier** | **Cat. number** |
| 5-fluorouracil (5-FU) | Sigma | F6627 |
| 78c | Calbiochem | 538763 |
| 8-Bromo-ADP-Ribose | BioLog | B 051 |
| 8-Bromo-cADP-Ribose | SCBT | sc-201514 |
| ACK lysing buffer | Thermo Fisher Scientific | A1049201 |
| Anti-Biotin MicroBeads | Miltenyi Biotec | 130-090-485 |
| BD Cytofix/Cytoperm Fixation/Permeabilization kit | BD Biosciences | 554714 |
| BrdU | Thermo Fisher Scientific | B23151 |
| DAPI | Molecular Probes | D1306 |
| Doxycycline hyclate | Applichem | APP A2951,0025 |
| DPBS 1x | Thermo Fisher Scientific | 14190-094 |
| Fetal bovine serum | Thermo Fisher Scientific | 1E+07 |
| Fixation/Permeabilization Concentrate | eBioscience | 00-5123-43 |
| Fixation/Permeabilization Diluent | eBioscience | 00-5223-56 |
| Fluo-8 AM | Abcam | ab142773 |
| HBSS buffer, $Ca^{2+}$+, $Mg^{2+}$+ | Thermo Fisher Scientific | 1E+07 |
| Horse serum | VWR | S0910-500 |
| Indo-1 AM | Thermo Fisher Scientific | I1223 |
| LS-columns | Miltenyi Biotec | 130-042-401 |
| MitoTracker Green FM | M7514 | Thermo Fisher Scientific |

| | | |
|---|---|---|
| normal rabbit serum | Abcam | ab7487 |
| PBS tablets | Sigma | P4417-100TAB |
| Permeabilization Buffer (10X) | eBioscience | 00-8333-56 |
| Pluronic F-127 | Thermo Fisher Scientific | P6866 |
| polyinosic:polycytidilic acid (pIC) | Tocris | 4287 |
| rabbit anti-mouse antiplatelet serum | Accurate Chemical and Scientific Corporation | WAK-AIA31440; |
| recombinant human FGF1 | Peprotech | 100-17A |
| recombinant murine IGF2 | R&D Systems | 792-MG-050 |
| recombinant murine SCF | PeproTech | 250-03 |
| recombinant murine thrombopoietin (THPO) | Thermo Fisher Scientific | 34-8686-63 |
| StemSpan™ SFEM | STEMCELL Technologies | 9650 |
| CellGenix TM GMP SCGM | CellGenix | 20802-0500 |
| Ficoll-Paque PLUS | GE Healthcare | 17-1440-03 |
| CliniMACS® CD34 MicroBeads | Miltenyi Biotec | 200-070-100 |
| recombinant human FGF1 | Peprotech | Cat. # 100-17A |
| Recombinant human FLT3 ligand | Miltenyi | Cat. #130-096-474 |
| Recombinant human SCF | Miltenyi | Cat. 130-096-692 |
| Recombinant human IL-3 | Miltenyi | Cat. #130-093-909 |
| T-5224 | AmBeed | A132992 |
| PEG300 | | |
| Tween80 | | |
| Tetramethylrhodamine ethyl ester (TMRE) | BioVision | K238 |
| Thapsigargin | Sigma | T9033 |
| Verapamil hydrochloride | Sigma | V4629 |
| nicotinamide guanine dinucleotide sodium salt (NGD) | Sigma | N5131-5MG |

Supplementary Figure 1

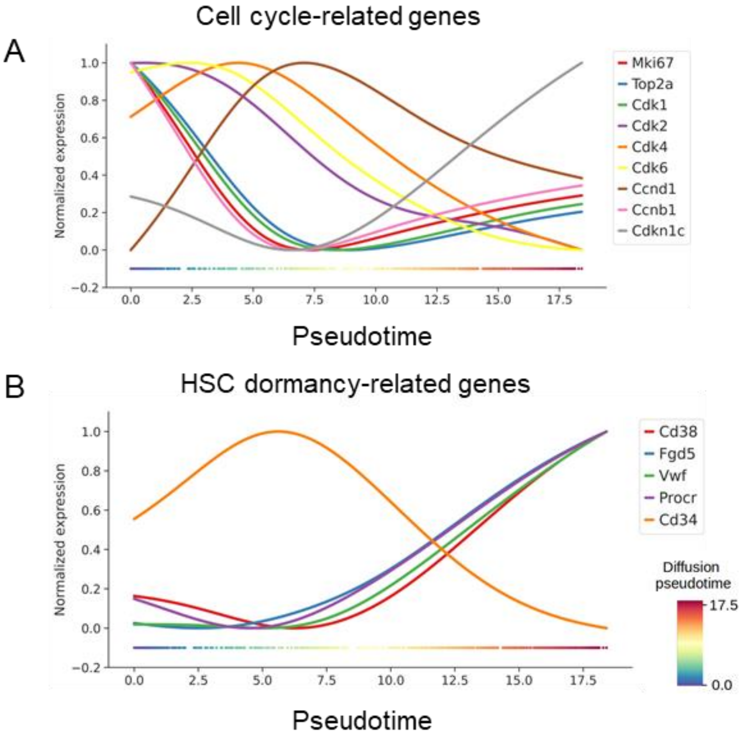


**Suppl. Figure 1. Single cell transcriptome analysis of HSCs.**

(A) Aligned kinetic curves for selected cell cycle-related genes along pseudotime. (B) Aligned kinetic curves for selected HSCs' dormancy related genes along pseudotime.

Supplementary Figure 2

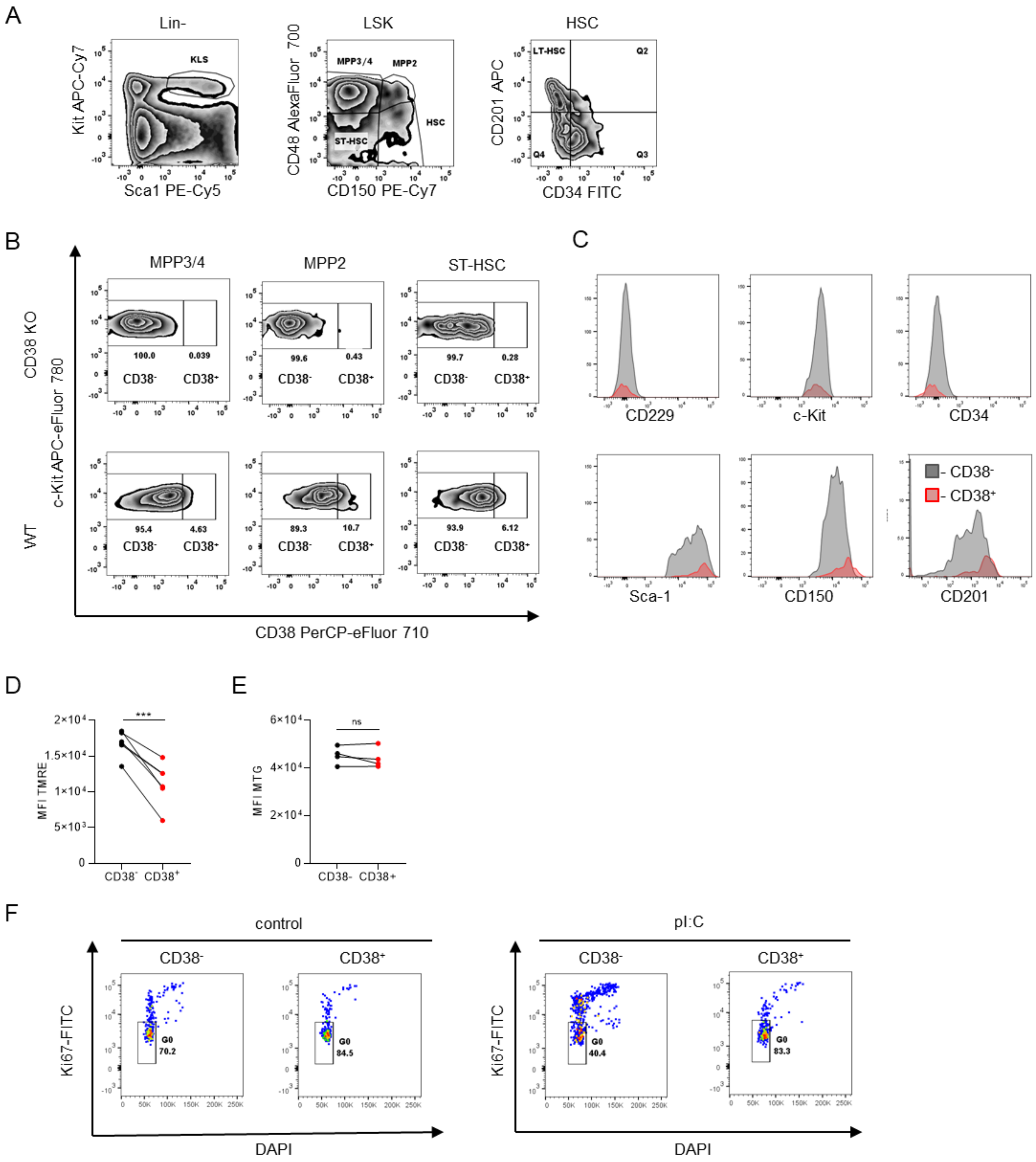


50

**Suppl. Figure 2. Characterization of CD38$^{+}$ and CD38$^{-}$ HSPCs.**

(A) Gating strategy for analysis of murine HSPCs. (B) Flow cytometry analysis of CD38 expression in HSPCs compartment (MPP3/4: Lin$^{-}$ Sca-1$^{+}$ Kit$^{+}$ (LSK) CD48$^{+}$ CD150$^{-}$, MPP2: LSK CD48$^{+}$ CD150$^{+}$, ST-HSCs: LSK CD48$^{-}$ CD150$^{-}$, HSCs: LSK CD48$^{-}$ CD150$^{+}$). HSPCs from CD38KO were used as negative control. (C) FACS analysis of defined markers surface expression on CD38$^{-}$ and CD38$^{+}$ HSCs. (D) HSC mitochondrial membrane potential analysis (n=5). (F) Analysis of mitochondrial mass in HSCs using MitoTracker Green (MTG) in the presence of verapamil (n=4). (G) Gating strategy for analysis of LT-HSCs in G0 phase of the cell cycle. P-values were calculated using a paired t-test, $^{**}p<0.01$, $^{***}p<0.001$.

### Supplementary Figure 3

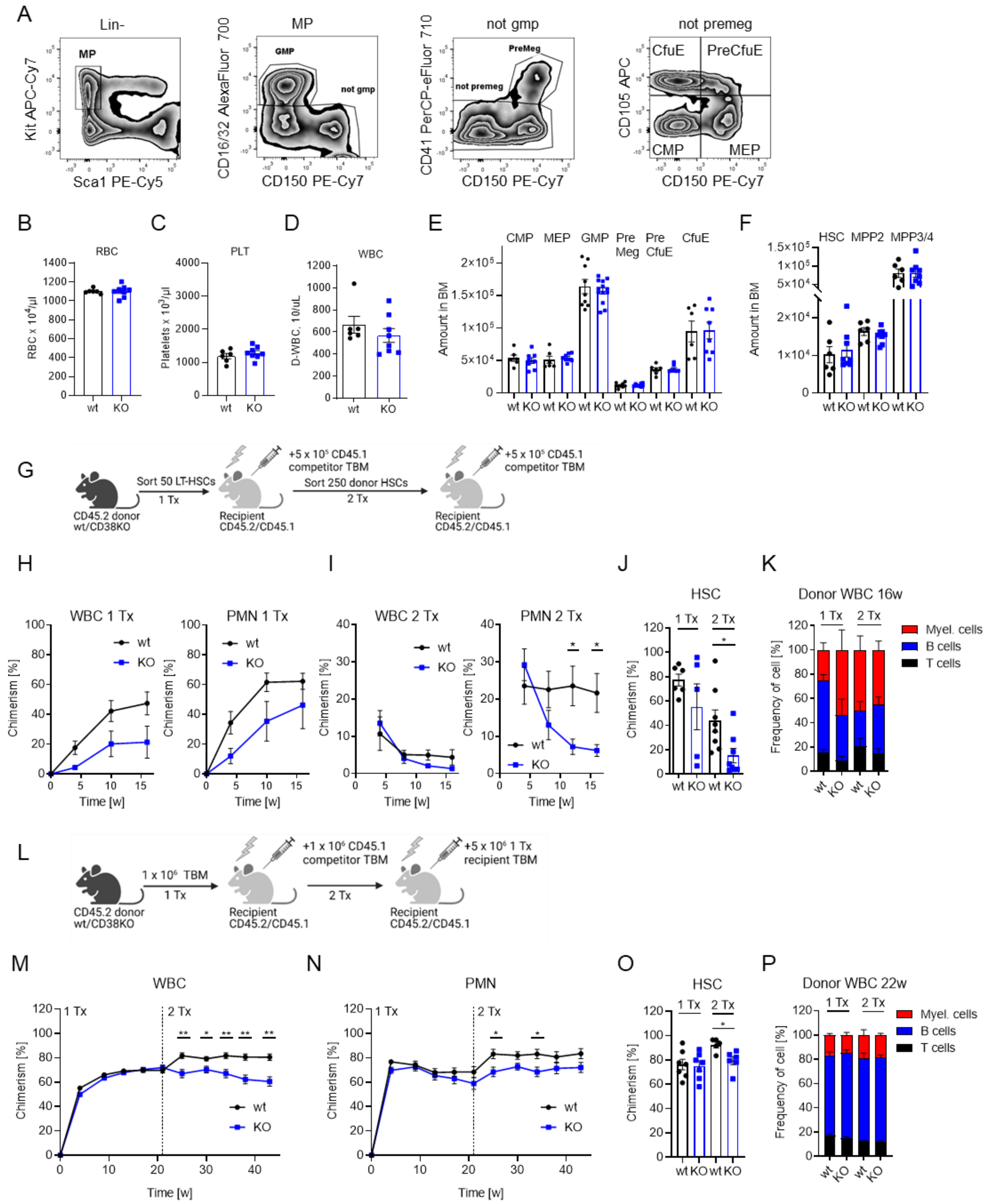

**Suppl. Figure 3. Functionality of LT-HSCs from CD38KO mice.**

(A) Gating strategy for analysis of restricted myeloid progenitors. (B) Number of RBC in peripheral blood of wt and CD38 KO (KO) mice. (C) Number of platelets (PLT) in peripheral blood of wt and CD38 KO mice. (D) Number of WBC in PB of wt and CD38 KO mice. (E) Number of restricted progenitors in bone marrow of wt and CD38KO mice. (F) Number of HSPCs in bone marrow of wt and CD38KO. (G) Experimental setup for transplantation of LT-HSCs from wt and CD38KO mice, 2 independent experiments, one representative experiment is shown. (H) Chimerism in donor-derived WBC and PMN ($Gr1^+$ $CD11b^+$) in peripheral blood (PB) of recipients after primary transplantation (n=5-6). (I) Chimerism in donor-derived WBC and PMN in peripheral blood (PB) of recipients after secondary transplantation (n=7). (J) Chimerism in donor-derived HSCs in bone marrow 16 weeks after transplantation. (K) Frequency of myeloid, B, and T cells in donor-derived WBCs 16 weeks after transplantation. (L) Experimental setup for transplantation of TBM from wt and CD38KO mice. (M) Chimerism in donor-derived WBC and PMN (N) ($Gr1^+$ $CD11b^+$) in peripheral blood (PB) of recipients after primary transplantation (n=5-6). (O) Chimerism in donor-derived HSCs in bone marrow 16 weeks after secondary transplantation (n=7). (P) Frequency of myeloid, B, and T cells in donor-derived WBCs 22 weeks after transplantation. P-values were calculated using Mann-Whitney U-test, *$p<0.05$, **$p<0.01$.

Supplementary Figure 4

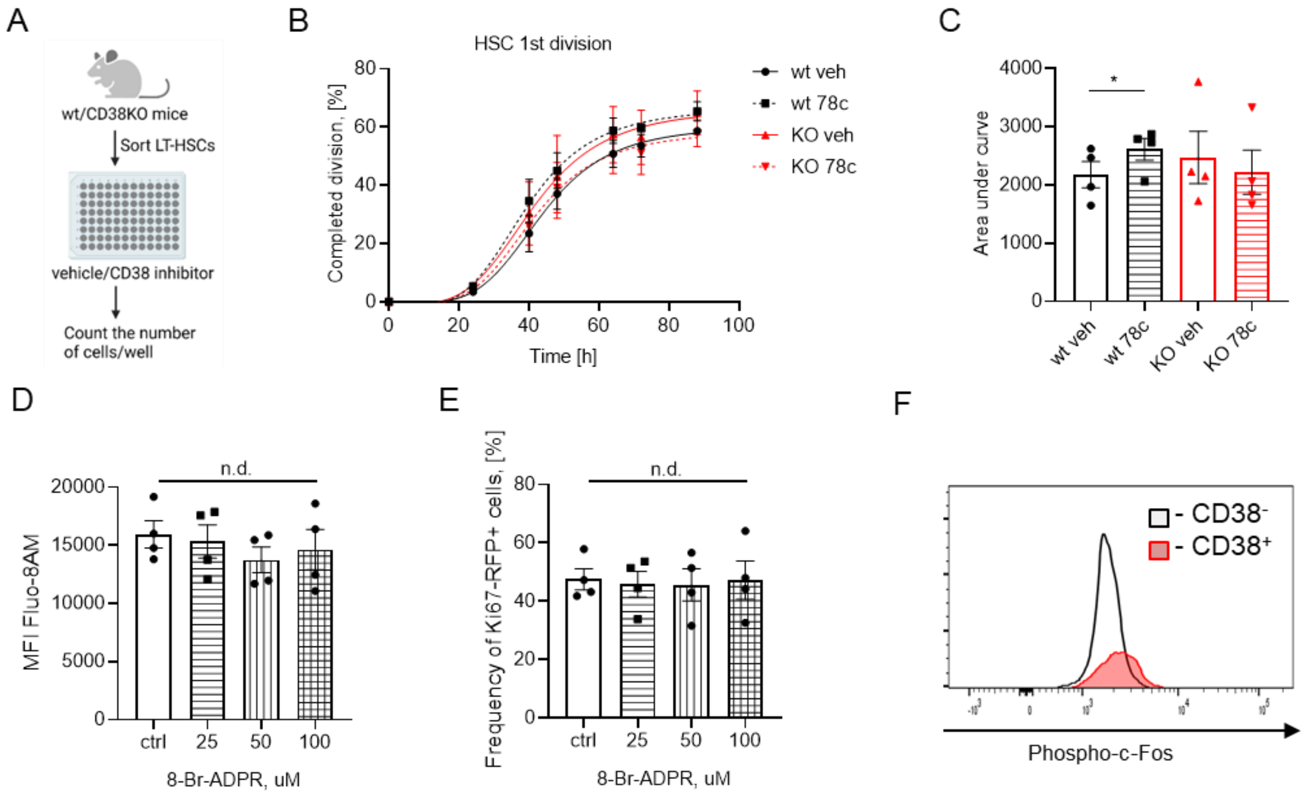


**Suppl. Figure 4. Regulation on LT-HSC division.**

(A) Setup for single cell division tracing experiment. Single LT-HSCs from wt and CD38KO were sorted and cultured in liquid media with or without 78c. (B) Frequency of LT-HSCs that had completed the first division during incubation time is presented (4 independent experiments). (C) Quantification of AUC for (B). Paired t-test, *p<0.05. (D) LSK from Ki67$^{ki/ki}$ RFP reporter mice were sorted and cultured for 24 h in the presence of ADPR antagonist (Br-ADPR). Relative $[Ca^{2+}]_c$ concentration in HSCs treated with Br-ADPR, (n=4). (E) Frequency of Ki67-RFP$^{+}$ HSCs 24 h after treatment with Br-cADPR, (n=4). (F) Representative plot of intracellular p-c-Fos in CD38$^{-}$ and CD38$^{+}$ HSCs. Multiple-group comparisons were performed using Brown-Forsythe and Welch ANOVA followed by Dunnett's T3 multiple comparison tests.

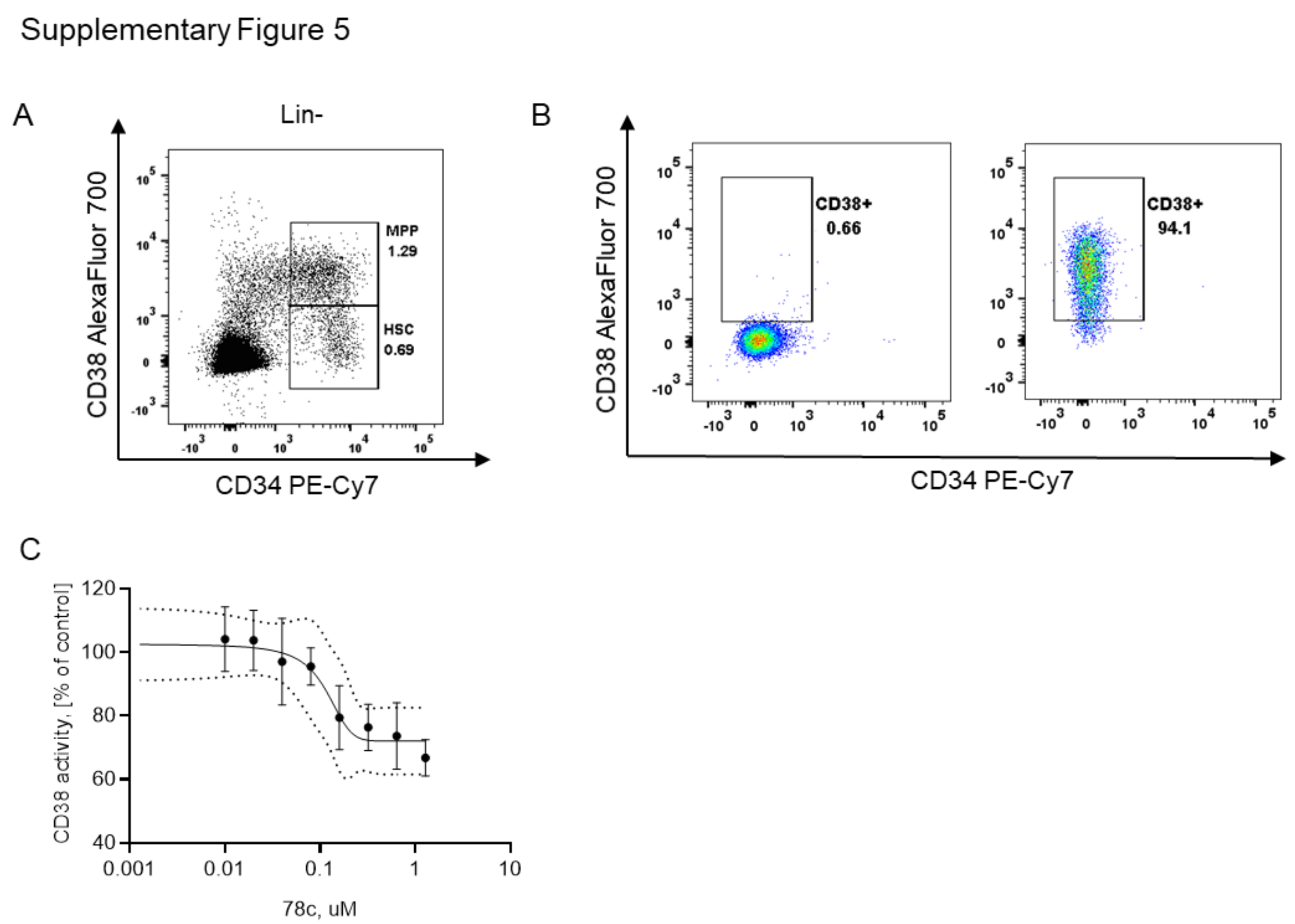


**Suppl. Figure 5. Expression of CD38 in human cells.**

(A) Gating strategy for isolation of human HSCs. (B) Surface expression of CD38 on MOLM-13. Left – isotype ctrl, right – anti-CD38 antibody. (C) Dose response curve for cyclase activity of MOLM-13 lysates in response to CD38 inhibitor, n=3. Sigmoidal standard curve was interpolated, 95% confidence interval is shown.

### References

1. Fornes O, et al. JASPAR 2020: update of the open-access database of transcription factor binding profiles. *Nucleic Acids Res.* 2020;48(D1):D87-D92.
2. de Oliveira GC, et al. Measuring CD38 Hydrolase and Cyclase Activities: 1,N(6)-Ethenonicotinamide Adenine Dinucleotide (epsilon-NAD) and Nicotinamide Guanine Dinucleotide (NGD) Fluorescence-based Methods. *Bio Protoc.* 2018;8(14).
